# Lipid-Driven Alignment and Binding of p7 Dimers in Early Oligomer Assembly

**DOI:** 10.1101/2025.06.13.659606

**Authors:** Oluwatoyin Campbell, Dina Dahhan, Viviana Monje

## Abstract

Proteins engage in interactions with lipid membranes to facilitate important cellular processes that form the basis of healthy or sick biological conditions. Transmembrane proteins such as ion channels are often made up of bound monomers that engage in specific contacts within the bilayer. However, molecular mechanistic information related to how these channel structures form is scarce. Understanding the role of lipids in driving this process have the potential to close knowledge gaps regarding the assembly behavior of oligomeric proteins which are often clinical targets for disease treatment. Using the hepatitis C virus p7 hexamer as a case study, this work focuses on characterizing the interactions that dictate the beginnings of dimerization using molecular dynamics simulations. Results comparing dimers formed in aqueous solution to those at the surface of a lipid membrane model reveal that protein-lipid interactions are critical in aiding the proper alignment and binding of inter-protein residues. Hydrophobic protein-lipid interactions and hydrogen bonding of key residues to phosphatidylcholine and phosphatidylinositol membrane lipids drive the characteristic inter-protein helix interactions that underlie p7 oligomerization. This increases favorable binding between hydrophobic protein residues, particularly for the first helix of the p7 monomers. This study provides evidence that membrane lipids are a necessary and dynamic factor that contributes to appropriate binding and association of proteins for channel formation within cellular membranes.

## I. INTRODUCTION

Protein and lipid interactions are fundamental contributors to cellular processes in health and disease. Lipid structural and compositional diversity in cell membranes provide a platform for protein interactions which underlie their function [1]. This involves changes to the structure of both proteins and lipids, which leads to the alteration of the surrounding lipid content near the protein and formation of sub-domains of different physical properties in the membrane [2-6]. These spatial and conformational changes are indicators of the influence of biochemical interactions and the cellular environment on protein structural dynamics and activity [1,7].

A class of proteins that has been well studied is transmembrane proteins, which contribute to cellular mechanisms of various types. Ion channels are a sub-class of membrane-embedded proteins that depend on voltage or ligand stimuli to move ions through lipid membranes [8-10]. Specific lipids within the immediate membrane environment can act as ligands to prompt opening and closure of the channel in different processes [11]. These structures are proposed to initiate by association of individual monomers, with folding happening alongside interactions between proteins and lipids to facilitate further aggregation during the assembly process [12]. However, the mechanistic details on how channels form are not well understood, preventing fundamental understanding of the process, especially within the context of maligned cell function. A promising tool to unravel these details is molecular dynamics (MD) simulations, which is based on statistical mechanics calculations of interaction forces. This technique has proven to be advantageous for modeling and highlighting protein-protein and protein-lipid interactions that drive mechanisms of biomolecules at the microscale [13-15].

In the context of disease, viroporins are ion channels that enable virus production by contributing to various steps in the viral life cycle [16-18]. In hepatitis C, the p7 viral ion channel features a complex structure that consists of intricately, intertwined monomers made up of 63 amino acids (Fig. 1A) [19]. The p7 protein stands out because of its important role in facilitating continuous virus production. Its influence on viral assembly, lipid droplet metabolism and localization near the lipid bodies have been well established, with these functions enabled through collaborations with other hepatitis C virus (HCV) proteins [20-22]. However, despite these key contributions, the assembly process of the p7 channel remains yet unclear.

**Figure 1.**
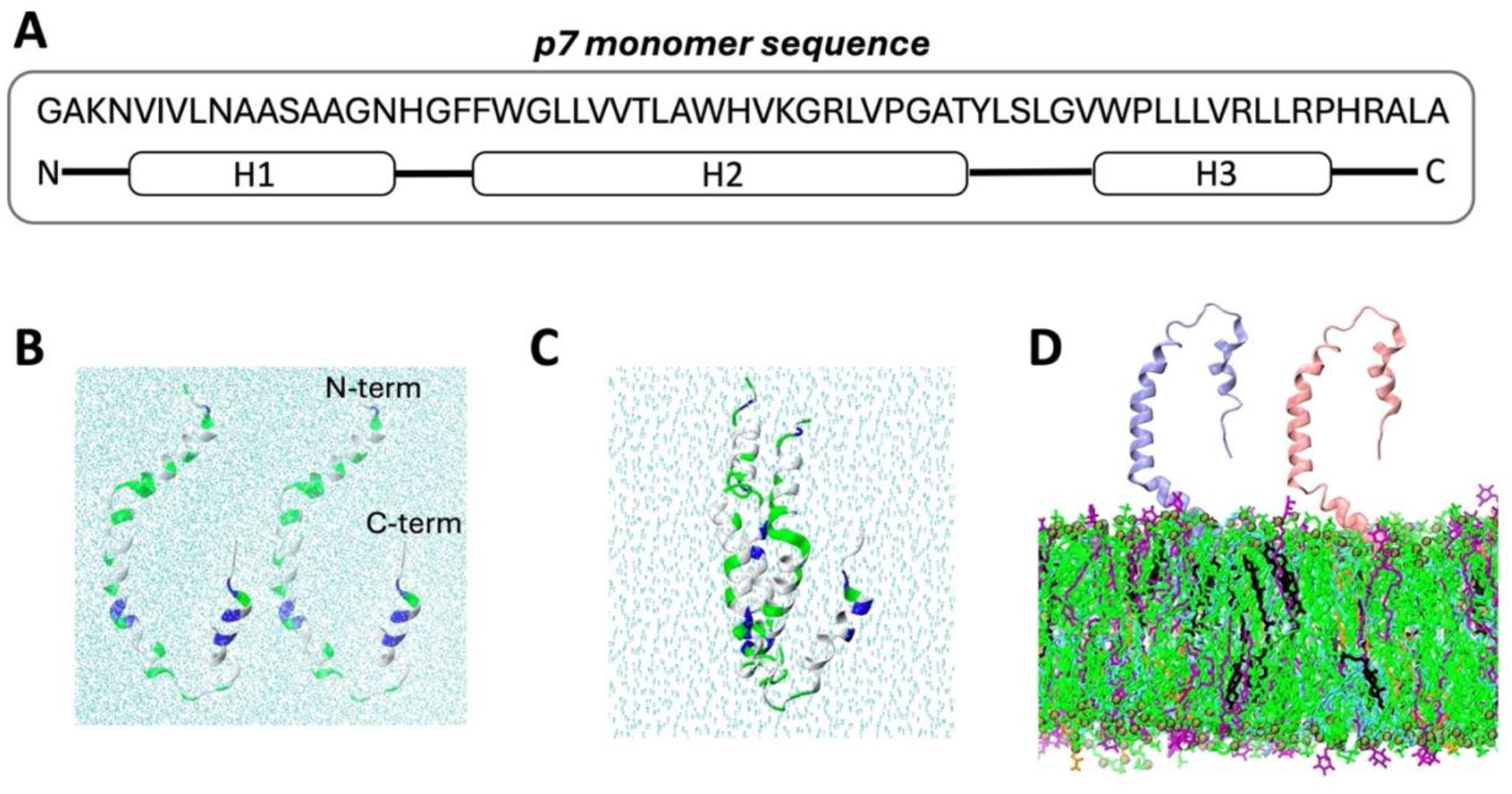
Initial simulation coordinates. **A)** p7 amino acids and their corresponding position within the protein. **B)** Two p7 monomers separated (*Sep* model) and **C)** extracted from the hexameric structure PDBID 2M6X (*Bound* model). Water atoms shown with sky blue. Proteins, with N and C termini indicated, are shown with cartoon representation, with non-polar residues in white, polar in green and cationic in blue. **D)** Two monomers initially attached by N-termini to the surface of a complex membrane (*Surface* model) containing DOPC (in green), DPPE (in blue), POPI (in purple), Cholesterol (in black) and DOPS (in orange) (55:21:11:9:4 mol%). Monomers are in the initial positioning seen in replica 1. In snapshots, water and ions are omitted for clarity.

Experimental investigations have revealed the role of anionic lipids in sustaining channel structure and membrane permeabilization activity [23,24]. The mechanism of p7 has been suggested to rely on the interplay between lipids and proteins that alter structural and mechanical properties of the local lipid environment. Molecular studies on p7 have focused on the role of specific protein residues in sustaining the spatial conformation of p7 monomers and hexameric channels within the membrane. Arg33 and Arg35 interact with negatively-charged lipid headgroups to sustain the transmembrane structure of p7 [24], cholesterol prompts a cascade of interactions that position His17 properly within the channel pore [6] and Tyr42 and Tyr45 sustain the kinks in the monomer membrane-embedded structure by increasing charged lipids presence nearby [25]. Our initial simulation studies on the interaction mechanisms of p7 with model bilayers explore the insertion of a single p7 monomer in the presence and absence of phosphatidylserine (PS) lipids. PS led to deeper insertion due to stronger hydrophobic and electrostatic interactions of the membrane with the first helix of the protein, which prompted increased disorder of the surrounding lipid environment [26]. This work seeks to address the role of lipids in mediating protein-protein interactions to facilitate the dimerization of p7 in the early stages of channel formation.

There are still unanswered questions about the mechanism of protein-protein association during transmembrane ion-channel formation at the molecular level. Fundamental understanding of these can aid in the design and development of therapeutics that target the action of homo-oligomeric membrane channels during viral disease pathogenesis. We hypothesize that the process is mediated by lipids at the surface of the membrane. To probe the importance of membrane lipids in p7 dimerization and its complex channel formation, we run microsecond-long classical MD simulations of two p7 monomers in water and near a membrane model for the endoplasmic reticulum (ER). Results consistently show that dimerization patterns of p7 heavily rely on hydrophobic and polar interactions with zwitterionic PC and anionic PI membrane lipids, specifically for protein membrane adsorption. These interactions provide a strong foundation that fixes p7 monomers in place to facilitate important hydrophobic interactions between them, especially in the first helix of its N-terminus. This work gives a detailed account of the function of apolar and charged lipids in mediating oligomeric protein structures.

## II. METHODS

To probe the effect of lipid composition on dimerization of p7 monomers, three different models were simulated: (i) the monomers initially separated in water (*Sep* model), (ii) a control system containing two monomers in water in the experimentally determined channel conformation from PDBID 2M6X (*Bound* model), and (iii) the monomers initially separated on the surface of an anionic-charged membrane (*Surface* model). Figures 1B-D illustrates the setup of each model, while Fig. S1 in the Supplemental Material [27] shows the different starting positions of the monomers in the *Surface* model. The membrane present in the *Surface* systems is based on the ER [23,28], containing 600 lipids per leaflet with 55% dioleoyl-phosphatidylcholine (DOPC), 21% dipalmitoyl-phosphatidylethanolamine (DPPE), 11% 1-palmitoyl-2-oleoyl-phosphatidylinositol (POPI), 9% cholesterol and 4% dioleoyl-phosphatidylserine (DOPS) lipid molecules. See Table S1 of the Supplemental Material [27] for relevant details of all simulated systems, including the respective number of replicas and simulation length. A total of 6.4 μs of trajectories using classical all-atom MD simulations were collected for this study.

The *Bound* and *Sep* models were constructed by placing two monomers from the 2M6X PDB channel structure in an aqueous solution box using CHARMM-GUI *Solution Builder* [29-31]. After the 4-step relaxation protocol generated by the builder, each model was run for 400 ns. For the *Surface* model, the bilayers were set up using the CHARMM-GUI *Membrane Builder* [29-33]. After completing the 6-step relaxation protocol generated by CHARMM-GUI, each membrane was equilibrated for 200 ns. Afterwards, an in-house bash script was used to merge the equilibrated membrane and monomers coordinates and run for 1 μs each. All systems were fully hydrated and neutralized with 0.150 molar KCl.

The CHARMM36m forcefield [34,35] which has been extensively validated with resulting structure of protein and lipids from experiments was applied in all simulations. The TIP3 model [36] was used to represent water molecules in the systems. The GROMACS package [37] was used to perform molecular dynamics on each model system. A timestep of 2 fs was chosen, which allows appropriate capture of vibrations of covalent bonded hydrogens constrained by the LINCS algorithm [38]. Electrostatic and van der Waals (vdW) non-bonded interactions were calculated with the Particle Mesh Edwards [39] and Verlet [40] algorithms. vdW interactions were modeled based on a Lennard-Jones potential force-switch function with a soft cutoff starting from 1.0 nm. Integration was carried out with the leap-frog integrator [41]. Temperature and pressure were set to 315.15K and 1 bar to capture body temperature and atmospheric pressure conditions. The Nose-Hoover [42,43] and Parrinello-Rahman [44,45] thermostat and barostat were used, with coupling times of 1.0 and 5.0 ps respectively. Compressibility was set at 4.5e-5 bar^-1^, based on that of water. Groups containing protein, membrane and water atoms were coupled separately to the 315 K temperature bath, and pressure was controlled semi-isotropically.

Analysis of resulting data was performed with in-house scripts that feature built-in tools from GROMACS and VMD, and Python libraries such as Numpy, Pandas, and MDAnalysis [46,47]. Analysis was computed using block averaging with respective standard errors for the latter half of the trajectories, utilizing at least 200ns of trajectory. A distance cutoff of 14 Å was used to define contacts between atoms to ensure sufficient sampling of lipids and/or amino acids. A full description of key analysis can be found in the supporting information, and sample scripts are publicly available at https://github.com/monjegroup/p7-dimer. In interaction analyses, interhelical interactions are referred to in the sequence of the first, then second monomer, e.g. H3-H2 refers to helix 3 of monomer 1 interacting with helix 2 of monomer 2. Snapshots were rendered with VMD [48]; in all instances, water and ions are not shown for clarity.

## III. RESULTS

### A. Dimer conformational changes are mediated by membrane lipids

After simulations were completed, the structural and conformational changes within the monomers were first evaluated. The *Sep* and *Surface* model systems (refer to Figure 1), equilibrated within the first half of the respective trajectory, as seen from the root mean square deviation (RMSD) time series of monomer 1 and monomer 2 (referred to as p1 and p2 in Fig. S2A-B in the Supplemental Material [27]). Therefore, the last 200ns of the *Sep* models, and the last 500 ns of the *Surface* models were used to calculate averages and standard errors of all properties reported for each replica. The RMSD of both p7 monomers is greater in the *Sep* model compared to the *Surface* model (p1: 1.40 ± 0.03 nm versus 1.29 ± 0.03 nm; p2: 1.37 ± 0.04 nm versus 1.22 ± 0.03 nm), corresponding to more notable change in secondary structure for the proteins in water. To compare both structures to the monomer conformation in the channel, the RMSD was also calculated using the *Bound* coordinates as the reference (Fig. S2C in the Supplemental Material [27]). The *Bound* model acts as a reference, with a conformation that initiates as the coordinates extracted from PDBID 2M6X (Fig. 2A). The difference is lower for monomer 2 in the *Surface* model, indicating more similarity to the *Bound* configuration in the presence of lipids.

**Figure 2.**
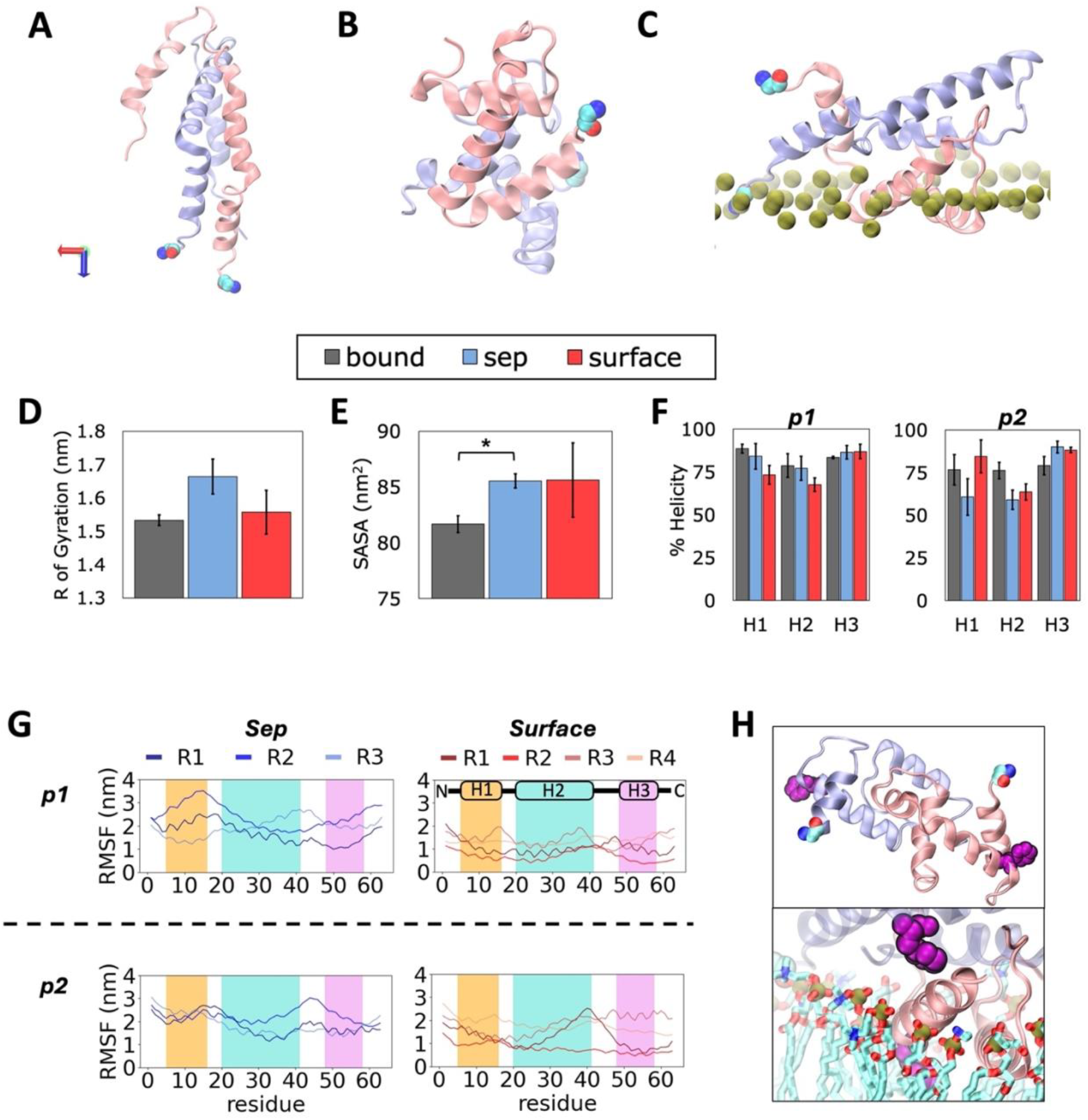
Protein conformational dynamics induced by presence of a lipid membrane. **A)** Initial conformation of dimers in the *Bound* reference model obtained from the p7 channel structure (available as PDB 2M6X). **B)** Representative final conformations of dimer structures formed in water (*Sep* model) and **C)** lipid (*Surface* model) environments. **D)** Average R_g_, **E)** SASA and **F)** protein % helicity in each model. **G)** RMSF of protein residues in *Sep* and *Surface* replicas. **H)** Snapshot of the Phe20 in helix 2 (H2), illustrating difference in localization in *Sep* and *Surface* models. Monomer 1 (p1) shown in pink, monomer 2 (p2) in ice-blue and N-terminus end indicated with van der Waal representation. Phosphorus atoms are shown in green and lipid tails in cyan. In all snapshots, water and ions hidden for clarity. Error bars represent standard error across replicas, and “*” indicates significant difference in means (p < 0.05).

The representative final conformations in each model can be seen in Figure 2; p1 and p2 monomers are colored in pink and ice-blue, respectively. Compared to the *Bound* model (Fig. 2A), the *Sep* model exhibits a globular dimeric structure (Fig. 2B), while the monomers stretch out as they interact with the membrane interface in the *Surface* model (Fig. 2C). To determine the depth of entry of each protein residue, the z-component of the distance between its center of mass (COM) and the average position of the lipid phosphorus atoms was calculated over the last 500 ns of trajectory. In general, the monomers remained above the membrane hydrophobic phase. In p1, residues 46-54 (Gly-Val-Trp-Pro-Leu-Leu-Leu-Val-Arg) are found closest to the membrane lipids, while p2 has Gly1 and Ala63 (N- and C-termini respectively) closest to the membrane (Fig. S3 in the Supplemental Material [27]).

When compared with the *Bound* model, the Radius of gyration (R_g_) of the *Surface* model appears lower, implying a more compact dimer structure, while that of the *Sep* model is larger (Fig. 2D). However, the difference between *Sep* and *Bound* is not statistically significant (p > 0.05). It is worth noting that the *Sep* model has a statistically greater solvent accessible surface area (SASA) than the *Bound*, while the *Surface* model does not (Fig. 2E). This implies that the exposure of residues to water is greater in the *Sep* model, and may act as a deterrent against proper alignment of monomers. To probe how much of the secondary structure is retained in the monomers after inter-protein contacts, the helical percentage of each of the three helices in each monomer was measured in the last half of the trajectories and reported in Figure 2F. These helices, referred to as H1, H2 and H3 hereafter, correspond to residues Val5 to Asn16, Phe20 to Thr41 and Trp48 to Pro58 in the p7 structure [19] (see. Fig 1A). The helical structure of p1 in the *Sep* model is comparable to the *Bound* reference, while that of *Surface* p1 loses a small amount of helicity in H1 and H2. In contrast, when compared with the *Bound* structure, *Sep* p2 loses while *Surface* p2 gains helicity in H2 and H3, respectively. The average helical content of H1 of the *Surface* model is greater than both *Bound* and *Sep* models.

To learn more about the dynamics of individual residues within the dimers, the root mean squared fluctuation (RMSF) of p7 residues was analyzed over the entire trajectory of each model replica and reported in Figure 2G. In general, the RMSF is greater for *Sep* replicas than the *Surface* replicas. Replicas do not exhibit the exact same RMSF profile, which is expected; yet, similar trends of peaks and minima can be identified for each model, particularly for residues found at the end of H1, which show higher fluctuation in *Sep* model versus the *Surface* model. *Sep* H2 helix has decreasing RMSF as opposed to *Surface* H2 that shows increasing RMSF along the helix, while H3 displays no sustained trend, implying its lower involvement during dimer formation in water or on the membrane surface. This is expected as H3 are the farthest apart between the monomers in the reference structure (Fig. 2A). The peaks and minima in the *Surface* model correspond to residues that interact with water or membrane lipids, respectively. Take for example, Phe20 at the start of H2 in each monomer (Fig. 2H), it faces the water phase in the *Sep* model (top panel) but it points to the membrane interior and interacts with the other monomer in the *Surface* model (bottom panel). These results show the role of membrane lipids in the formation of protein-protein contacts and conformational changes, which facilitate dimer formation.

### B. Lipids contribute to the residue alignment of p7 dimers

To shed more light on the inter-protein alignment behavior of p7 in the studied models, an inter-residue contact analysis was carried out with a cutoff of 14 Å. The fraction of time protein contacts occurred during the equilibrated trajectories in the *Sep* and *Surface* models was compared against those of the initial 50 ns of the *Bound* reference structure. Contact maps for the full set of replicas of each model is shown in Fig. S4 in the Supplemental Material [27]. The conserved contacts between monomers across all replicas of the *Bound* model is indicated with black rectangles. Figure 3A highlights main trends across models. In the *Sep* model, conserved target interactions observed in the *Bound* reference which involve 1-to-1 (i.e. H1-H1, H2-H2 and H3-H3 contacts) and H3-H2 helix contacts are reduced, with those in the H1-H1 region completely missing. In comparison, the *Surface* model containing the anionic membrane lipids exhibits contacts in the two critical regions that correspond to the reference state. Therefore, though full dimerization was not seen, the lipid membrane clearly facilitates proper H1-H1 and H3-H2 interactions.

**Figure 3.**
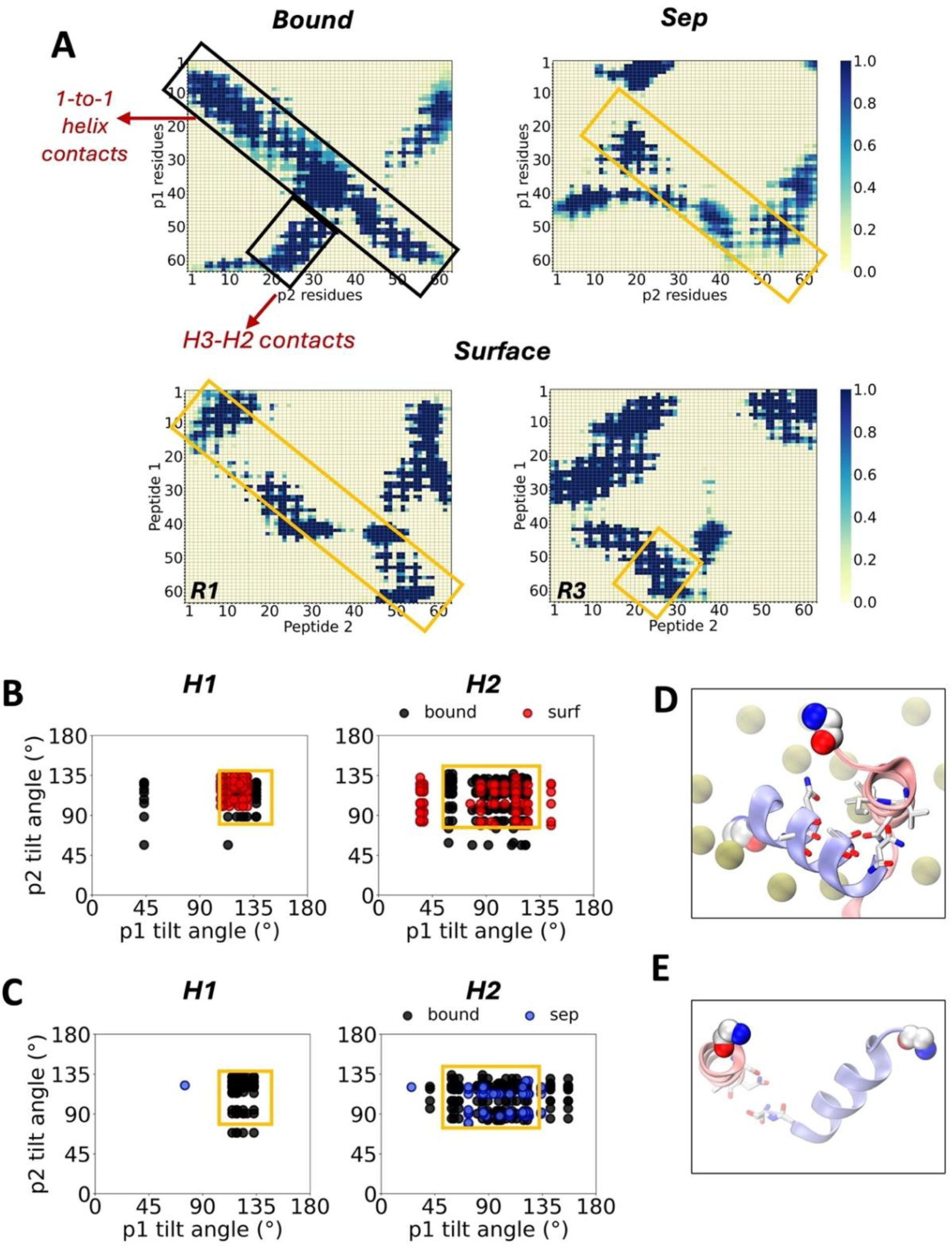
Residue contact comparison with reference dimer structure. **A)** Contact maps of residues, based on a cutoff of 14 Å. *Surface* replicas 1 (R1) and 3 (R3) shown in bottom panel. **B)** Tilt angle conformational landscapes of 1-to-1 contacting helices in p1 and p2 in representative *Surface* and **C)** *Sep* replicas. **D)** Snapshots showing helix 1 contacts in *Surface* and **E)** *Sep* representative replicas. Contacts and tilt angles indicate trends during second half of system trajectories. Target contacts conserved across all *Bound* reference model replicas (1-to-1 inter-helical contacts and helix 3-helix 2 contacts) indicated with black rectangles. Contact regions formed in *Sep* and *Surface* models indicated with yellow rectangles. Carbon, oxygen and nitrogen atoms shown in white, red and blue respectively.

Next, the conformational landscape of the interacting residues involved in the 1-to-1 helix alignments were further evaluated. For residues with a contact occupancy fraction of at least 0.9, the tilt angle of the side chain was measured with respect to the vector defined between alpha carbons of residues with low flexibility as determined from the trajectory movie (see Table S2 of the Supplemental Material which includes references [30,35] therein). These reference alpha carbons were as follows: Val7 and Ala14 for H1, Gly22 and Lys33 for H2 and Pro49 and Leu56 for H3. The full set of results for *Sep* and *Surface* replicas can be seen in Figs. S5 and S6 in the Supplemental Material [27], respectively. There is a conserved region across all replicas of the *Bound* model in the H1 and H2 conformation maps (indicated in yellow). The same region is observed in the corresponding maps for H2 in the *Sep* and *Surface* models, while that of H1 is only reproduced in the *Surface* model (Fig. 3B and 3C). *Surface* representative snapshots show contacts facilitated by both polar and hydrophobic residues in H1 as the N-terminus of p2 anchors both helices to the membrane (Fig. 3D). In contrasts, H1 helices in the dimer in water are consistently apart from one another (Fig. 3E), suggesting membrane lipids promote H1-H1 interactions.

When comparing contact and conformational maps of all *Surface* replicas, a unique behavior can be seen in replica 4, which has an absence of points in the conserved regions of both the contact and 2D tilt angle maps (Figs. S4 and S6 in the Supplemental Material [27]). SASA and R_g_ measurements of this replica in Fig. S7A-B in the Supplemental Material [27] show it deviates from the behavior observed in the other *Surface* replicas in that the monomers are further apart (see also Fig. S7C in the Supplemental Material [27] vs Fig. 2C). This may result due to the orientation of the N-terminus of p2 that points outwards to the aqueous phase, as p1 moves towards the leaflet and fully interacts with membrane lipids during initial steps of the simulation (Fig. S7D in the Supplemental Material [27]); this behavior is absent in other replicas.

### C. Protein binding interactions are energetically favored in the presence of lipids

The structural and conformational analyses above show that the monomers align in a manner more similar to the *Bound* reference structure in the presence of lipids. To identify key interactions that drive such a distinct dimerization mechanism in presence of lipids, the energetics of solvation, non-bonded interaction, and free energy of inter-monomer binding are reported in Figure 4. The average solvation energy is lower for the *Surface* model compared to the *Sep* model (Fig. 4A), implying the cost of monomer dissociation is lower and more favorable in the presence of lipids. The reference structure from the channel exhibited the highest solvation energy, showing that the energy that must be overcome to detach the binding monomers in the *Bound* model is greater than *Sep* and *Surface* models.

**Figure 4.**
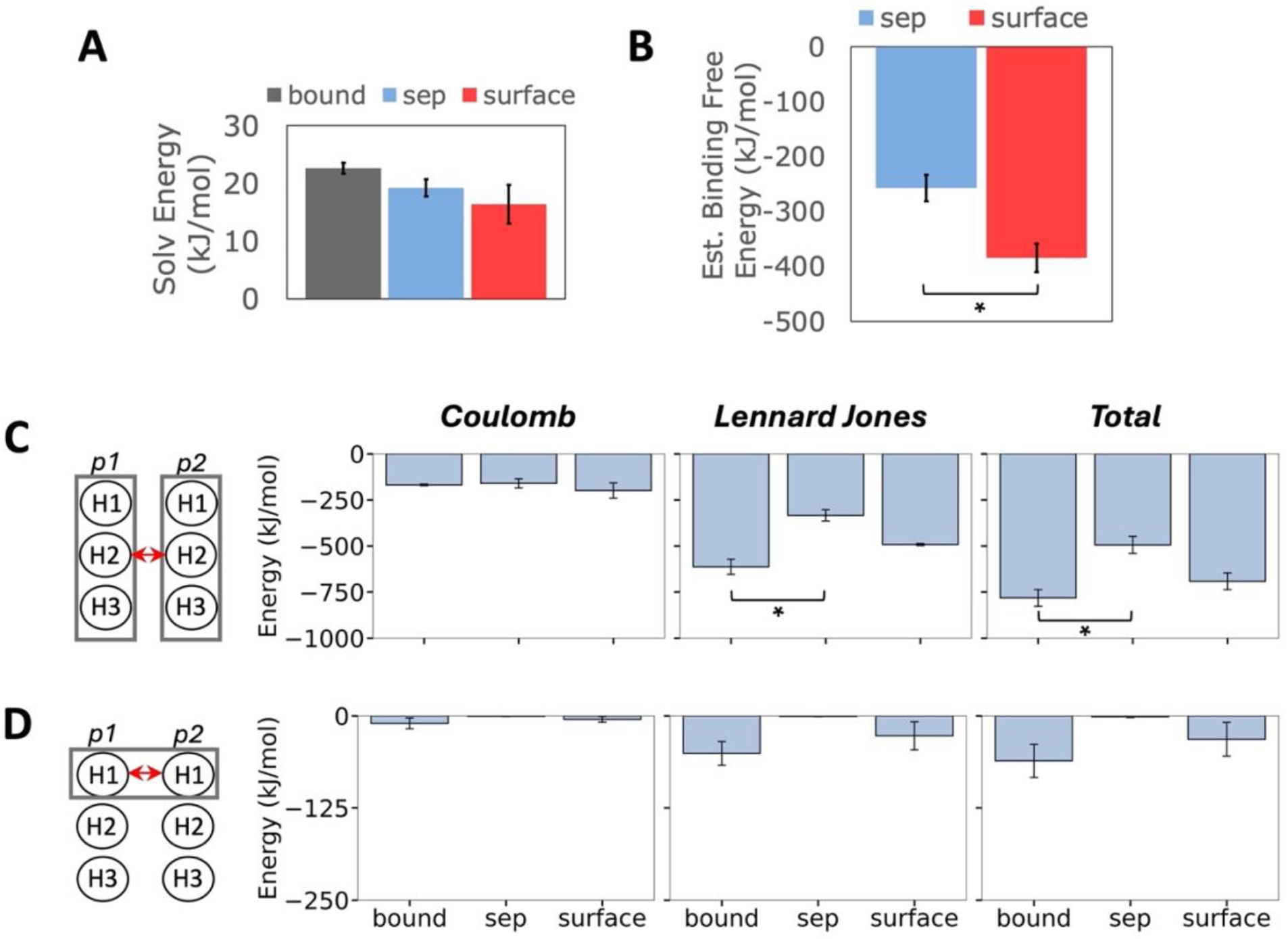
Energetics of protein interactions. **A)** Solvation energy of the formed dimer in each model. **B)** Estimate of the protein binding free energy per model, computed with the Molecular Mechanics with Generalized Born and Surface Area (MM-GBSA) approach. **C)** Coulomb (electrostatic), Lennard-Jones (hydrophobic) and total interaction energy of monomer 1 and monomer 2 and **D)** helix 1 and helix 1 in each model. These energies are shown in the first, second and third column respectively. Error bars represent standard error across replicas, and “*” indicates significant difference in means (p < 0.05). Statistics computed exclude replica 4.

The binding free energy was estimated using the Molecular Mechanics Generalized Born Surface Area (MM-GBSA) method, further described in the supporting information. These estimates take into account the solvation energy described by the Generalized-Born equation and the interaction energy of in-vacuum binding; using the gmx_MMPBSA software [49], the free energy difference with respect to the bound state was determined. Figure 4B shows that the binding free energy is statistically more favorable in the *Surface* model compared to the *Sep* model, supporting the hypothesis that membrane lipids are important drivers for inter-protein binding and formation of p7 dimers.

To evaluate the contribution from non-bonded interactions, the electrostatic (Coulomb) and hydrophobic (Lennard Jones) energies were determined using the *gmx rerun* command in GROMACS. There is little difference in electrostatic contributions across models. In contrast, hydrophobic interactions are markedly larger in the *Surface* model, leading to statistically more favorable interactions between the monomers at the membrane surface (Fig. 4C). The total H1-H1 non-bonded interaction energy in the *Surface* model is comparable to the *Bound* reference, while that of *Sep* model is effectively zero (Fig. 4D). This shows more favorable binding between monomers at the membrane leaflet, which led to more accurate structure and conformation of the *Surface* dimer structure. This suggests that hydrophobic interactions play an important role in driving protein alignment for the formation of p7 dimers.

Non-zero estimates of the non-bonded energy for other helical interactions are also shown in Fig. S8 in the Supplemental Material [27]. As in the full monomer case, the greatest differences between the *Sep* and *Surface* models arise to favor hydrophobic interactions. Of note, H2-H2 interactions are less favorable in both *Sep* and *Surface* models than in the reference structure, while H3-H3 interactions are overrepresented in both compared to the reference structure. Lastly, H2-H3 interactions are miniscule, while H3-H2 interaction favorability falls short in both *Sep* and *Surface* models in comparison to the *Bound* structure. This difference in H2-H3 vs H3-H2 interactions is understandable, as though the sequences of the two monomers are identical, each helix and residue sees a different chemical environment based on individual distance from other molecules in the system.

### D. Lipids enhance anticorrelated protein-lipid motions

To determine specific protein-lipid interactions during dimer assembly, the contact frequency of these structures was analyzed during the last 500 ns of trajectory, using 14 Å as the cutoff distance. The results for the *Surface* replicas with the most similar inter-protein contacts compared to the *Bound* structure are shown in Figure 5A, replica 1 showcases 1-to-1 helix alignment, and replica 3 that of H3-H2 alignment. Both replicas feature notable presence of DOPC, POPI and DPPE contacts with p1, the closest monomer to the membrane. Cholesterol (indicated in red) interacts mostly with H1 and H2 residues in p1 and the N-terminus of p2, all of which enter deeper into the membrane interior compared to others (see Fig. S9 in the Supplemental Material [27]), and stabilize the protein dimerization process. DOPS contacts show up only in the case of H3-H2 alignment as it interacts with residues at the N-terminus and H3 of p1 (indicated in green). Our results suggest this interaction may be needed to stabilize H3-H2 alignment for accurate dimer formation.

**Figure 5.**
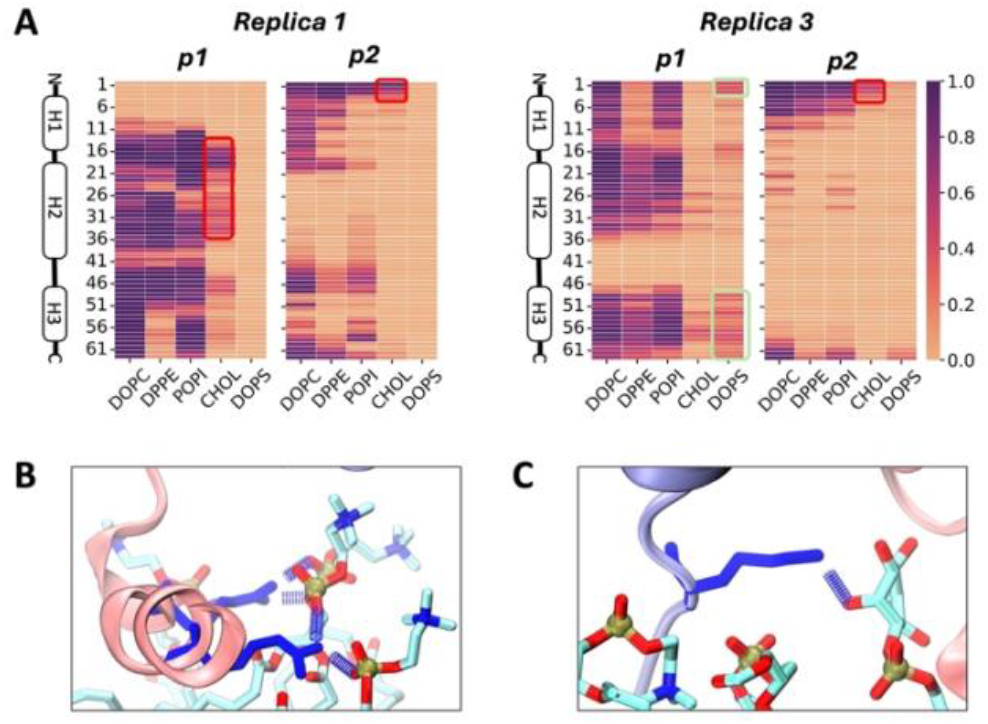
Protein-lipid interactions in the *Surface* model. **A)** Frequency of contacts between each protein residue and lipid species in the second half of the trajectory, based on a cutoff of 14 Å. Results shown are representative *Surface* replicas which had the best alignment of the dimers. Cholesterol and DOPS contacts highlighted with red and light green rectangles respectively. Snapshots illustrating some important hydrogen bonds: **B)** Arg residues in helix 3 of p1 with DOPC lipids, and **C)** Lys residue in N-terminus of p2 with a POPI lipid. Arg and Lys residues shown in dark blue licorice representation. Lipid carbon, oxygen, nitrogen and phosphorus atoms are shown in cyan, red, blue and gold respectively.

The hydrogen bonding frequency between protein residues and lipids was also calculated for the last 500ns of trajectory in the *Surface* model, based on a distance and angle cutoff of 3.2 nm and 30°, respectively. The representative replicas show hydrogen bonds involving arginine residues in H3 and lysine at the tip of H1 with DOPC and POPI lipids (Fig. 5B and 5C). Once again, DOPS notably enhances H3-H2 alignment shown in replica 3, with p1 forming well-sustained hydrogen bonds with Lys3, Arg57 and Arg60 (see Fig. S10 in the Supplemental Material [27]). In replica 1, additional H-bonds formed with the 15-21 (Gly-Asn-His-Gly-Phe-Phe-Trp) sequence in the first loop after H1, and Trp30, His31 and Arg35 in H2 may be aiding the formation of proper H1-H1, H2-H2 and H3-H3 contacts (see Fig. S10 in the Supplemental Material [27]). The N-termini H-bonds strengthen monomer-lipid interactions to sustain H1-H1 alignment between the monomers.

Dynamic cross correlations (DCC) analysis was also conducted to investigate correlated behavior in protein-lipid dynamics using an adaptation of the algorithm reported in Tekpinar et al. [50]. This involved tracking the displacement of interacting protein alpha carbons and lipid phosphate atoms, and after which the time average of the dot product of individual carbon-phosphate combinations is computed. To minimize empty selections during the calculation, protein-lipid contacts were selected based in a cutoff of 20 Å. Positive correlations indicate unidirectional movement of protein-lipid pairs, while negative correlations indicate movement in opposite directions.

Figure 6 shows key results; the two *Surface* replicas with the greatest similarity to the *Bound* reference model stand out by displaying anti-correlated motions. When 1-to-1 helix alignment is present, the motions of residues 11-56 in p1 are anticorrelated with all 5 lipids, motions of residues 1-4 and 46-50 in p2 are anticorrelated with DPPE, and motions of residues 41-47 in p2 are anticorrelated with cholesterol (Fig. 6A). On the other hand, when H3-H2 contacts are present, motions of residues 32-39 in p1 are anticorrelated with the top 4 lipids, motions of residues 14-39 in p1 are anticorrelated with DOPS, and motions of residues 1-16 in p2 are anticorrelated with DPPE and DOPS (Fig. 6B). This is in contrast with the fully positive correlation between all residue-lipid combinations observed for the replicas that did not show target inter-protein binding (see Fig. S11 in the Supplemental Material [27]). These results imply that anticorrelated protein-lipid movement may be a marker for protein dimer assembly.

**Figure 6.**
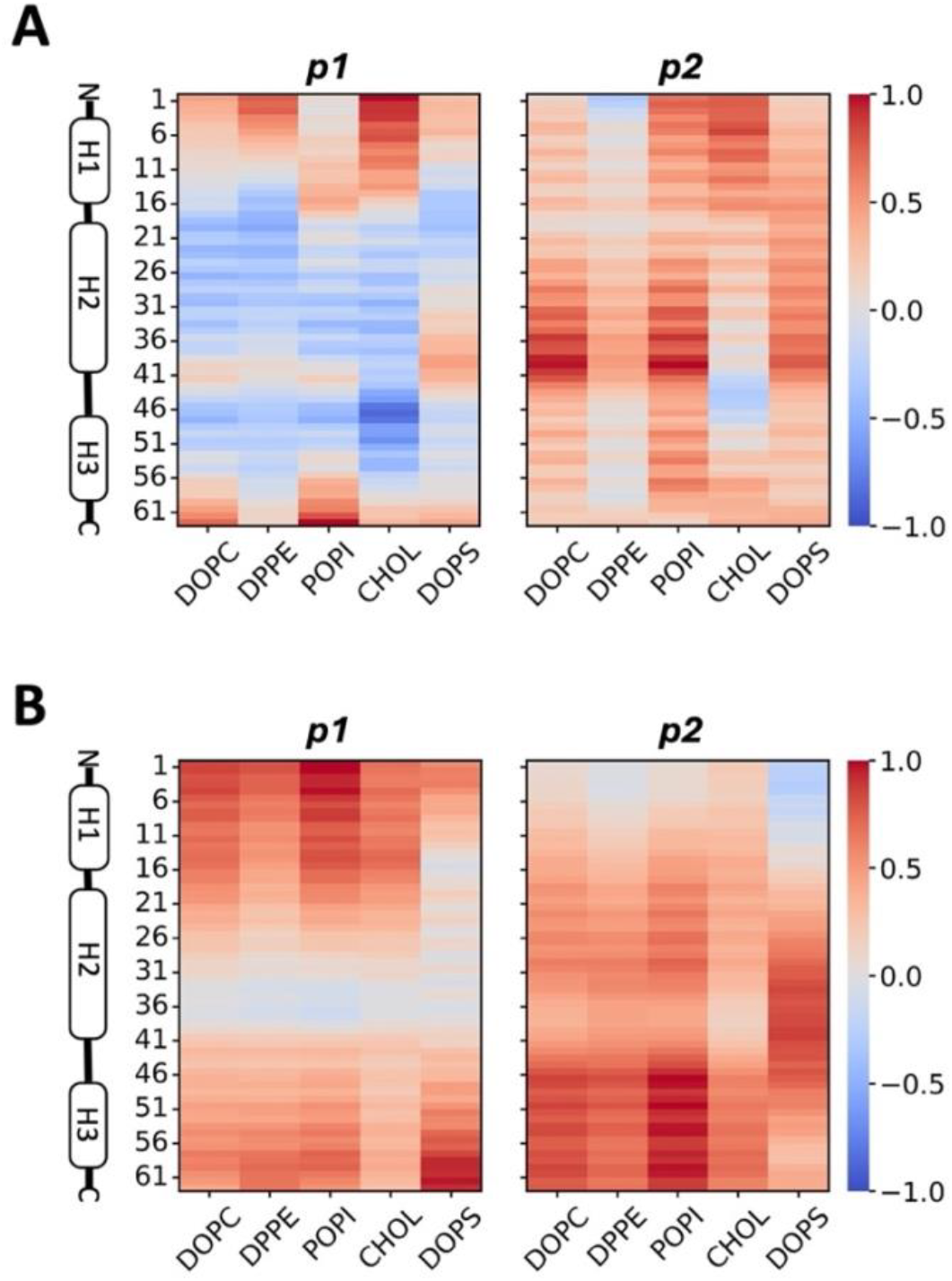
Correlation of protein and lipid motions in the *Surface* model. **A)** Map of dynamic cross correlations (DCC) of each monomer with each contacting lipid species in the replica 1 and **B)** replica 3 during the entire simulation trajectory. Negative correlation is represented in blue, and positive correlation in red. Contact is defined within a cutoff of 20 Å.

## IV. DISCUSSION

Protein assembly mechanisms of transmembrane homo-oligomeric channels remain underexplained for the most part. This stems from existing challenges in examining the dynamic changes of membrane protein structures at high resolution. Vesicles and lipid detergents used in experimental investigations are often limited to simple mixtures of membrane lipids compared to more complex models of the diverse lipidome of cells [51]. Techniques such as NMR, Xray crystallography and cryo-electron microscopy cannot capture detailed dynamics seen below the microsecond timescale, and cannot capture the specificity of lipid influence on these processes [52,53]. In studies of ion-channel forming proteins, efforts have focused on exploring pore characteristics within the membrane core [54-57], or the monomers already set in oligomeric form prior to interacting with the membrane interface [58]. Given the lack of understanding on how these structures oligomerize, we set out to explore the molecular mechanism by which a model ion channel forms in this work. Specifically, we investigated the mediatory role of membrane lipids in facilitating protein-protein interactions in the formation of p7 dimers from HCV. This study examines two identical monomers following the premise that ion channel assembly mechanism starts with the binding of two monomers into a dimer [12]. Our study analyzes long unbiased MD simulation trajectories of two p7 monomers in aqueous solution and at the membrane interface of a complex membrane model for the ER.

Evaluation of structural and conformational properties of the monomers emphasized differences in dimerization of the *Sep* model, which contains the monomers in water, and the *Surface* model, where the p7 monomers are near a model bilayer. Membrane lipids facilitate more compact dimer conformation, with increased helicity for H1 in one of the monomers versus its water counterpart. Similarly, the orientation of individual residue side chains shared increased similarity to the reference *Bound* state conformations for the dimer formed near the bilayer. H1, the first helix of p7, is amphipathic, featuring both polar and non-polar protein residues. Amphipathic proteins are well-characterized as an effective pore inducer within bacterial membranes in the context of antimicrobial activity [59]. The lower helicity in the amphipathic H1 of p7 in water is supported by studies of the aggregation mechanisms of membrane disrupting peptides from bacteria and parainfluenza, which report a negative impact of water on helical propensity [54,60].

The membrane acts as an anchor for the monomer in closer contact to the lipid headgroups, reducing residue fluctuation. The RMSF profile of the monomers showed a non-uniform adsorption behavior in both *Sep* and *Surface* models, as the residues with lowest and highest fluctuations were different in each model. Yet, differences were particularly distinct for residues at the end of H1 (orange region) in the presence of lipids, which had lower RMSF values. Inter-protein H1 contacts did not readily form in water, suggesting anchoring of at least one monomer by membrane lipids facilitates inter-helix alignment. In the fully formed hexameric channel, the first helix of p7 is found within the interior pore structure [19]. Given this eventual positioning, the amphipathic nature of this helix likely prompts dimer interactions to minimize hydrophobic strain as the protein inserts into the membrane interior.

The surrounding environment also influences protein conformations and interactions. This is demonstrated by changes in the 1-to-1 helix contacts, which led to different tilt angle combinations for individual residue side chains involved in the contact. The range of angles accessible by the residues in H1 of the nearest monomer to the membrane is more restricted than those of the other one; spanning 100^0^-145^0^ for H1 of p1, compared to 70^0^-135^0^ for H1 of p2. None of the *Sep* replicas access these conformations for correct H1-H1 alignment, while 3 out of 4 *Surface* replicas did. This implies that proper H1-H1 alignment of residue side chains between p7 monomers requires the assistance of lipid interactions.

The role of non-bonded interactions in facilitating dimer formation is non-trivial for inter-protein binding in both water and lipid environments. Though the contributions from electrostatic interactions were nearly equal in all model cases, differences were found in hydrophobic contributions, where Lennard Jones energy was larger and more favorable in the presence of membrane lipids. These results agree with conclusions of an experimental and computational study of the pore-forming channel Vp4 of hepatitis A virus, where mutations that reduced protein hydrophobicity led to attenuation of membrane entry and virus production [58]. Further analysis of the contributions of individual helical interactions identified H1-H1 interactions are only favored in the presence of the membrane. In comparison, H2-H2 and H3-H2 interactions were not observed in either aqueous or lipid environments, possibly because there are greater energetic barriers to be overcome to form target inter-helical contacts as the protein aggregates into higher order oligomers. Binding free energy and solvation energy estimates supports a role of the membrane in promoting favorable interactions between monomers during early p7 channel assembly. It has been shown that dimer formation of helix-containing proteins benefits from hydrophobic interactions provided by apolar bulky residues such as Leu, Ile and Trp [60,61]. The non-trivial role of electrostatic and polar interactions is also highlighted in driving and stabilizing the insertion of membrane-spanning homo-oligomeric structures [24,54]. Additionally, the *Surface* model did not induce deep entry of the dimeric p7 into the hydrophobic phase of the membrane compared to our previous monomer study. A replica exchange simulation study of a similar protein supports that the process of dimerization is more energetically costly than remaining in monomeric form [60], and this may explain this limited sampling of protein insertion. Therefore, to fully probe the oligomerization mechanism of p7, advanced sampling techniques may be needed.

Lipid dynamics have important mediatory roles in the formation of protein oligomers; lateral lipid reorganization allows the formation of membrane nano-domains that lower energetic costs for protein association [62]. In this study, systems that exhibit more accurate dimer assembly have a negative correlation between the dimer and surrounding lipids, which translates to an opposite direction of movement between the respective alpha carbon and phosphorus atoms in contact pairs. It has been noted that strong positive or negative correlations usually underscore important inter-protein communications during events such as binding of substrates to enzymes, which lead to functional changes in structure due to allostery effects [63]. Therefore, the presence of significant anti-correlation trends in this study indicates a greater involvement of lipids is needed for successful monomer alignment and assembly.

We also probed contact and hydrogen bonding patterns between p7 and neighboring lipid species to elucidate what type of interactions assist inter-protein binding at the membrane surface. The greater contact area of the monomer closer to the membrane and sustained contact of terminal residues of the monomer farther from the membrane seem to be key for proper peptide formation, with DOPC, POPI and DOPS playing important roles. In general, zwitterionic PC and anionic PI lipids contribute the most to p7 interactions, including sustained hydrogen bonds with cationic Arg and Lys residues. PS contributions are centered around pinning down H3 to the membrane surface. This is expected and in line with p7 structure, given that H3 contains the greatest presence of cationic Arg residues. The role of PS in facilitating protein-protein interactions during viral protein oligomerization is also reported in an experimental study of the VP40 matrix protein of the Marburg virus, where PS was highlighted as important for virus assembly and maturation [61].

Notably, our study also showed sustained cholesterol contacts with the p7 terminal ends and majority of H2 residues when there was 1-to-1 helix alignment. This implies a role of cholesterol in ensuring stable protein conformation that may drive oligomer formation, in agreement with results from a fluorescence spectroscopy study that shows cholesterol drives membrane penetration and oligomer formation of the amphiphilic viral fusion peptide of SARS-CoV [64]. Interestingly, this study also shows low and high cholesterol concentration modulate the two roles respectively. Hence, it is possible that local membrane transitions between cholesterol enriched and depleted regions are relevant for protein oligomerization. There is growing evidence in the literature that supports key roles of lipid composition in modulating protein-protein interactions on the membrane interface and its interior, which underscores a variety of cellular signaling processes in health and disease.

## V. CONCLUSIONS

The process of oligomeric assembly of ion-channel forming proteins is not fully understood, and there are currently little to no investigations that outline the detailed mechanism of this important biological phenomenon. In an effort to clarify this, this work probes the unique role of lipids in the initial step of oligomerization of the HCV p7 viroporin as a case study. This was conducted with MD simulations that examined binding and conformational changes of two p7 monomers in aqueous solution and at the interface of a complex ER membrane model.

Results establish that the presence of lipids prompts better protein alignment at the membrane interface, through the mechanism outlined in Figure 7. During first contact with the membrane, PC and PI lipids enable strong attachment of the monomers through hydrophobic and hydrogen bonding interactions with specific residues, favoring 1-to-1 helix interactions of H1, H2 and H3 from both monomers, as well as H3 and H2 contacts. These contacts stabilize conformations that allow favorable hydrophobic residue interactions, particularly in the case of H1-H1 alignment, as shown by protein helicity, inter-residue contacts, and residue side chain orientation detailed in this study. Taken together with our previous work [26], this study emphasizes the non-trivial role that specific membrane lipid species play in monomer membrane adsorption and dimerization.

**Figure 7.**
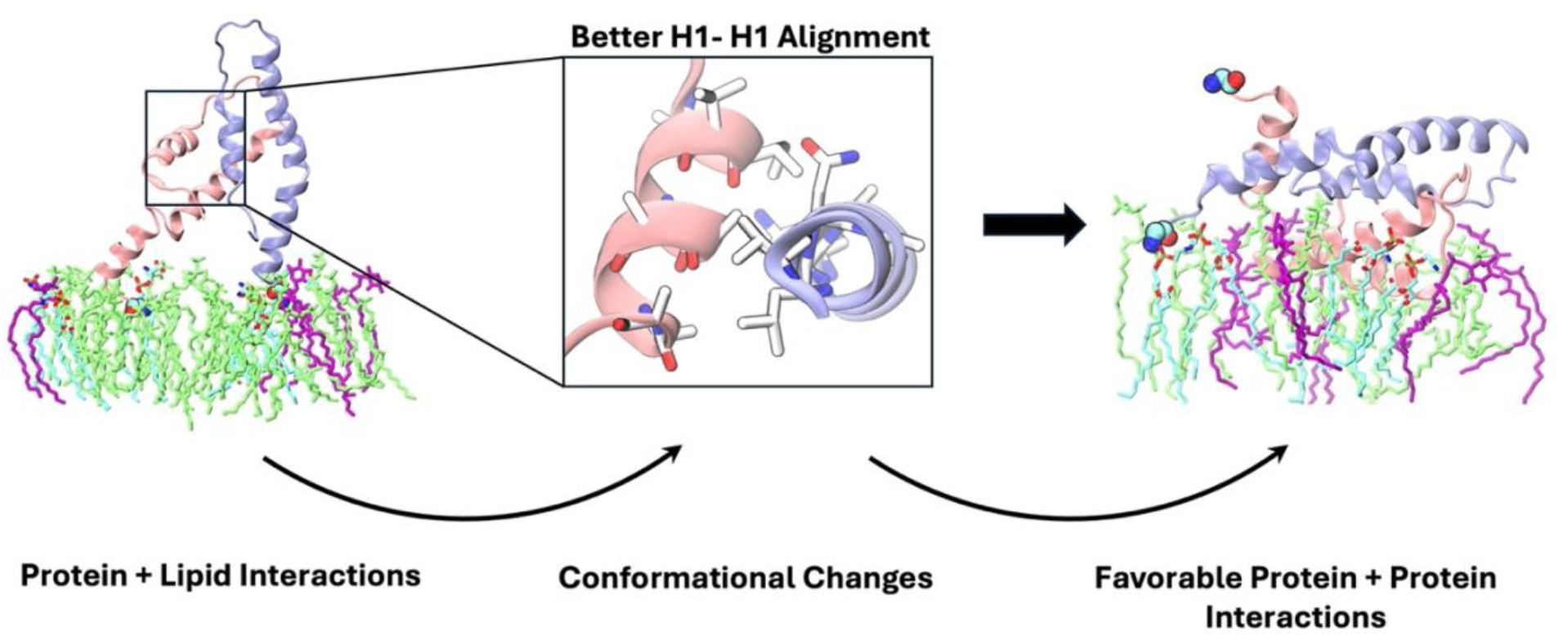
Suggested mechanism of p7 dimerization at the ER membrane surface, based on results from this study. DOPC and POPI lipids shown in light green and purple, helix 1 (H1) residues shown in white, oxygen atoms in red and nitrogen in blue. Monomers 1 and 2 differentiated with pink and ice-blue cartoon representation. Water and ions omitted for clarity during rendering.

This classical MD simulation study also highlights the energy barriers that exist in the process of dimerization. Future work should employ enhanced sampling techniques to provide an in-depth understanding of dynamics and interactions that orchestrate the assembly mechanism of homo-oligomeric channels composed of helical proteins. Such studies will help to elucidate the role of lipids in protein insertion and oligomerization for various viral proteins, as well as those involved in signaling pathways relevant to healthy and other diseased conditions that remain elusive.

## ACKNOWLEDGEMENTS

This work was performed on the University at Buffalo’s Center for Computational Research (Center for Computational Research, 2019). The authors thank the Graduate Women in Science for funding support of the research of O.C.’s through the Nell I. Mondy Fellowship. O.C. was also supported by the University at Buffalo Presidential Fellowship, and the National Institute of Health’s Initiative for Maximizing Student Development Training Grant T32 GM144920 awarded to Margarita L. Dubocovich (PI).

## AUTHOR CONTRIBUTIONS

O.C.: Investigation, Methodology, Data curation, Formal analysis, Validation, Writing – original draft. D.D.: Data curation, Analysis, Writing – original draft. V.M.: Conceptualization, Project administration, Supervision, Resources, Writing – review & editing.

## DATA AVAILABILITY STATEMENT

Final system coordinates and topology files are available at [10.5281/zenodo.15376633], simulation trajectories are available upon request. Python, Bash and TCL analysis scripts used to calculate helicity, frequency of residue-lipid contacts and hydrogen bonds, tilt angle conformational landscapes, dynamic cross correlations and binding free energies are publicly available at https://github.com/monjegroup/p7-dimer.

## APPENDIX: DESCRIPTION OF ANALYSIS

### A. Solvent accessible surface area (SASA)

SASA quantifies protein packing by taking a measure of the exposed surface area to surrounding solvent. Fully folded proteins with lower SASA values are generally considered more thermodynamically stable [65], making it a useful measure to draw conclusions about protein structure and equilibrium. It can be estimated with the Double Cubic Lattice Method (DCLM) algorithm from Eisenhaber et al, which uses 3 dimensional grids laid over the molecule to group spatially close atoms and indicate unobstructed atoms according to equation 1 [66,67].

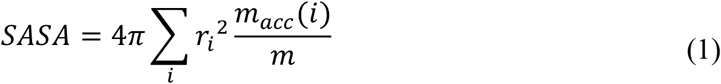

where *m* is the total number of dots that cover a unit sphere, *m*_*acc*_(*i*) is the number of dots on atom i not occluded by neighboring atoms, and *r*_*i*_ is the atomic radius. The summation is taken over all the atoms in the molecule.

### B. Radius of gyration (R_g_)

R_g_ can be used to gain further insight into the compactness of proteins; for example, a smaller R_g_ indicates a more compact molecule [68]. In simulations, it aids in determining the thermal stability of the system. *R*_*g*_ is defined as the square root of the mass-average of the radii components orthogonal to each axis. Over a time series, the evolution of this property can be computed according to equation 2:

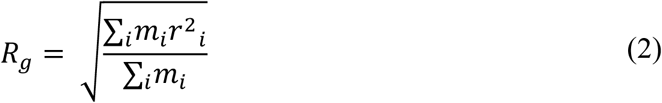

where *m*_*i*_ is the mass of atom i, and *r*_*i*_ is the position of atom i.

### C. Root mean square deviation and fluctuation (RMSD and RMSF)

RMSD quantifies shifts in internal coordinates of a structure by first aligning the atoms in each frame of the trajectory with a least-square fit using a user-specified reference frame and computing the distance between the two. RMSF quantifies the average over time of the RMSD to examine individual dynamics of residues or user-selected atoms; here it is presented for each protein residue. Both RMSD and RMSF can be used to determine equilibrium during a simulation trajectory and inform of sampling of specific conformational changes [69]. The RMSD is calculated according to equation 3:

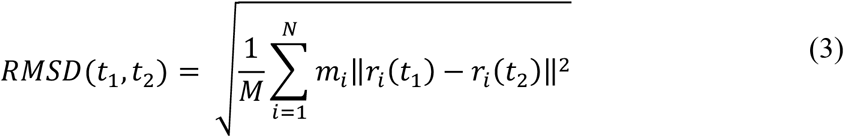

where 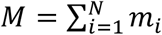 is the mass of atom i and *r*_*i*_(*t*) is the position of atom *i* at time *t*. The reference structure is found at time *t*_2_; If the initial structure is used as a reference, then *t*_2_ = 0. The RMSF is calculated as the time averaged value of the RMSD for each of the protein residues or user-selected atoms.

### D. Helicity

Helicity is a measure of the percentage of the amino acids in a user-defined selection that meet the biochemical criteria to be considered part of a helical structure. In this work, the criteria is based on the STRIDE algorithm, which uses a weighted contribution of hydrogen bonding patterns, energy and backbone torsion angle probabilities to define thresholds that fulfill the minimal conditions of secondary structure. Equations outlining the criteria are found in the methods section of [70].

### E. Residue-Lipid Hydrogen Bonds

Residue-lipid hydrogen bond analysis measures the strength of interactions between each residue and each type of phospholipid in the membrane by quantifying the frequency of hydrogen bonds formed over a chosen time slice. Hydrogen bonds are identified using the VMD *hbonds* plugin, where a bond is counted when the distance between the bond donor and the bond acceptor is less than 3.2 Å, and the angle between the donor to the hydrogen to the acceptor (angle D--H-A) is less than 30° [71].

### F. Tilt Angle Conformational Landscape

The conformational landscapes of protein residues give an understanding of how residue conformation changed in hand with sustained interactions between monomers. The algorithm to calculate this is as follows:

1. When a residue of the first monomer is within 14 Å of another in the second monomer, a contact is defined.
2. Strongly contacting residues are filtered based on if the frequency of contact in the time slice considered is greater than 0.9.
3. Reference vectors for each protein helix are defined using residues of low flexibility that hardly change in helical conformation.
4. Tilt angle vectors are defined as the vector connecting each residue’s alpha carbon and its terminal tail atom outlined in Table S1 in the Supplemental Material [27].
5. Using the vectors, the tilt angle of the residues that meet the criteria are plotted as coordinates on a 2D plane.

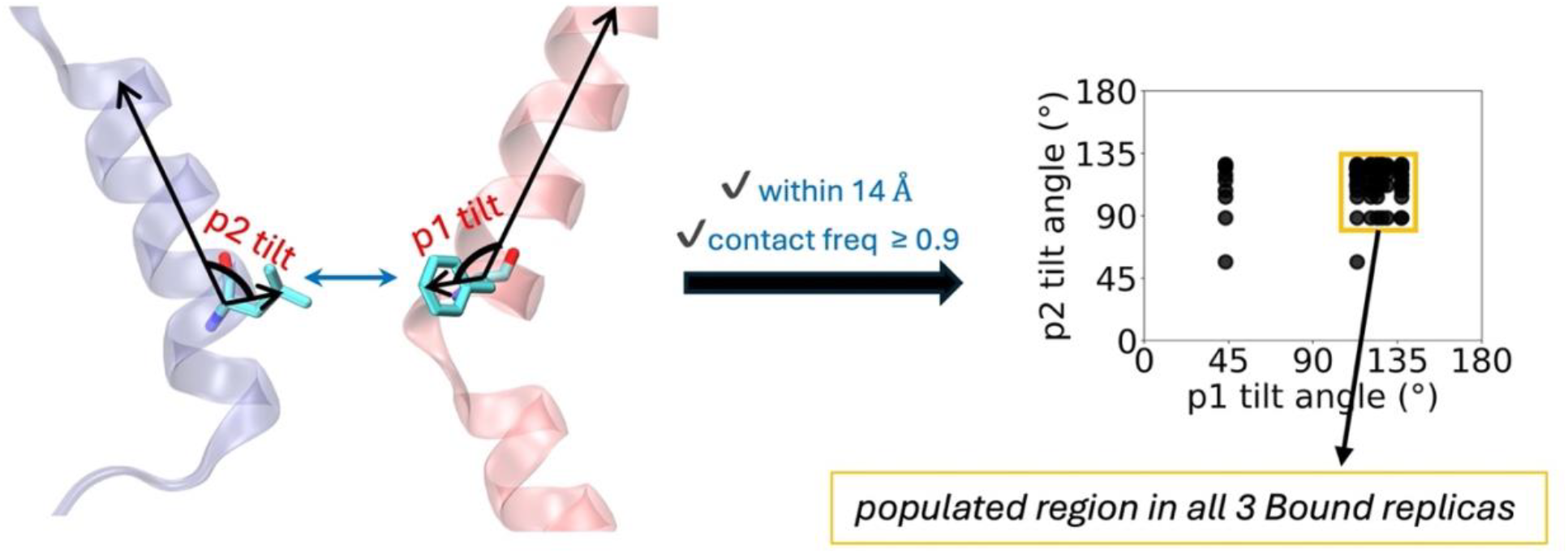

The figure above summarizes how the vectors and criteria were defined and provides guidance on how to interpret the measured tilt angles of a sample 2D plot.

### G. Dynamic Cross Correlation (DCC) Analysis

DCC is used to identify parts of a protein that experience correlated movement. This value can be positive, which indicates movement in the same direction, or negative, which indicates movement in opposite directions. The algorithm used in this work is adapted from Tekpinar et al [50], measuring the displacement of the protein alpha carbons relative to that of lipid headgroup phosphorus atoms within 20 Å of each other. The time average of the dot product between the positional vectors for each carbon-phosphorus combination was computed according to equation 4:

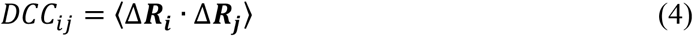

where Δ***R***_***x***_ = ***R***_***x***_ − ⟨***R***_***x***_⟩ is the positional difference vector of atom *i* with respect to its average position after an overall root mean square alignment procedure has been conducted.

### H. Binding Free Energy Estimates

Here, we use the Molecular Mechanics Generalized-Born Surface Area (MM-GBSA) method to estimate the energetic cost that must be overcome for protein binding. The binding free energy was calculated according to the following equations:

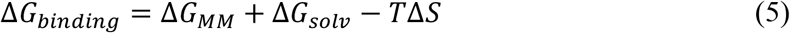

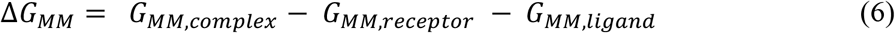

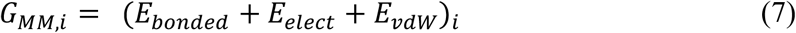

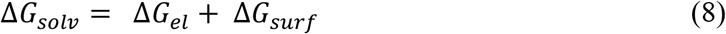

where Δ*G*_*MM*_ represents the difference in sum of the bonded (*E*_*bonded*_) and nonbonded (electrostatic, *E*_*elect*_ and van der Waals, *E*_*vdW*_) interaction energy of the complexed molecule and those of the individual components, which are usually termed the receptor and ligand respectively [72]. The Δ*G*_*solv*_ is the solvation energy, which estimates the polar contribution, Δ*G*_*el*_ with the Generalized-Born [73] equation, and the nonpolar contribution, Δ*G*_*surf*_ is based on a linear approximation of the solvent accessible surface area of the complex [49,74].

