## Supplemental Material for "Lipid-Driven Alignment and Binding of p7 Dimers in Early Oligomer Assembly"

**Table S1**. Description of systems studied. The composition of the bilayer in the *Surface* model was set as 55% DOPC, 21% DPPE, 11% POPI, 9% cholesterol and 4 % DOPS. Each membrane contained 1200 total lipids (600 per leaflet).

| model | replica | total atoms | water atom | box size (*x, y, z,* in nm) | sim. time (ns) |
|---|---|---|---|---|---|
| ***Bound (reference)*** | Rep1 | 212,693 | 210,297 | 13 x 13 x 13 | 400 |
| | Rep2 | 212,693 | 210,297 | 13 x 13 x 13 | 400 |
| | Rep3 | 212,693 | 210,297 | 13 x 13 x 13 | 400 |
| ***Sep (water)*** | Rep1 | 212,225 | 209,829 | 13 x 13 x 13 | 400 |
| | Rep2 | 214,349 | 211,953 | 13 x 13 x 13 | 400 |
| | Rep3 | 214,349 | 211,953 | 13 x 13 x 13 | 400 |
| ***Surface (membrane)*** | Rep1 | 528,710 | 371,586 | 18.8 x 18.8 x 14.6 | 1,000 |
| | Rep2 | 535,295 | 378,171 | 18.8 x 18.8 x 14.9 | 1,000 |
| | Rep3 | 534,764 | 377,640 | 18.8 x 18.8 x 14.9 | 1,000 |
| | Rep4 | 534,677 | 377,553 | 18.7 x 18.7 x 15.0 | 1,000 |

**Table S2**: List of protein residues and selected atoms to define the tilt angle vectors. Nomenclature corresponds to the C36m FF [see ref. 30,35 in the main manuscript]

| **Amino Acid** | **Atom names** | **Atom types** |
|---|---|---|
| GLY | HA1 | HB2 |
| ALA | CB | CT3 |
| VAL | CB | CT3 |
| LEU | CG | CT1 |
| ILE | CG1 | CT2 |
| PHE | CZ | CA |
| PRO | CG | CP2 |
| SER | OG | OH1 |
| THR | CB | CT1 |
| TYR | OH | OH1 |
| LYS | NZ | NH3 |
| ARG | CZ | C |
| HSD | CE1 | CPH2 |
| TRP | NE1 | NY |
| ASN | CG | CC |

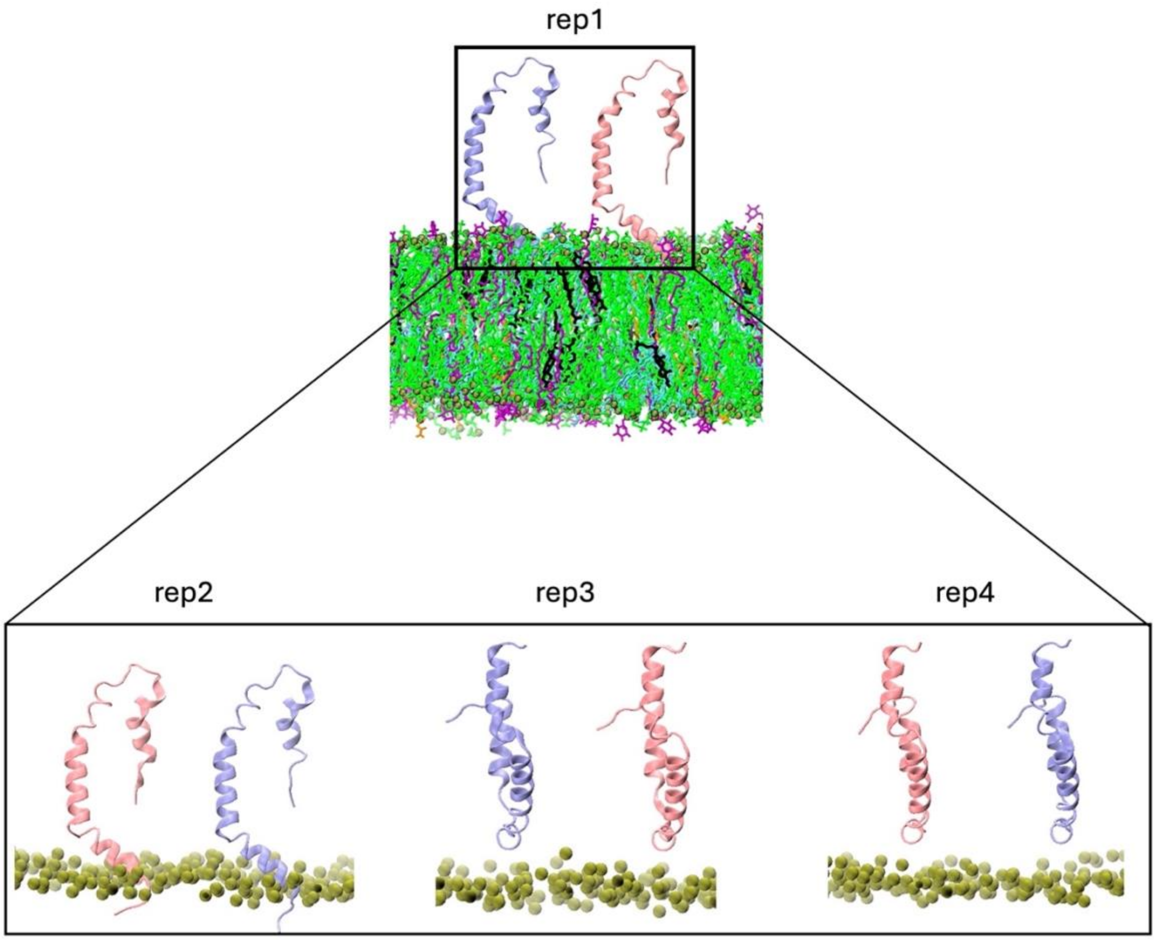


**Figure S1.** Initial positioning of monomers in replicas 1, 2, 3 and 4 of *Surface* model. Lipid phosphorus atoms shown in green to illustrate positioning with respect to membrane.

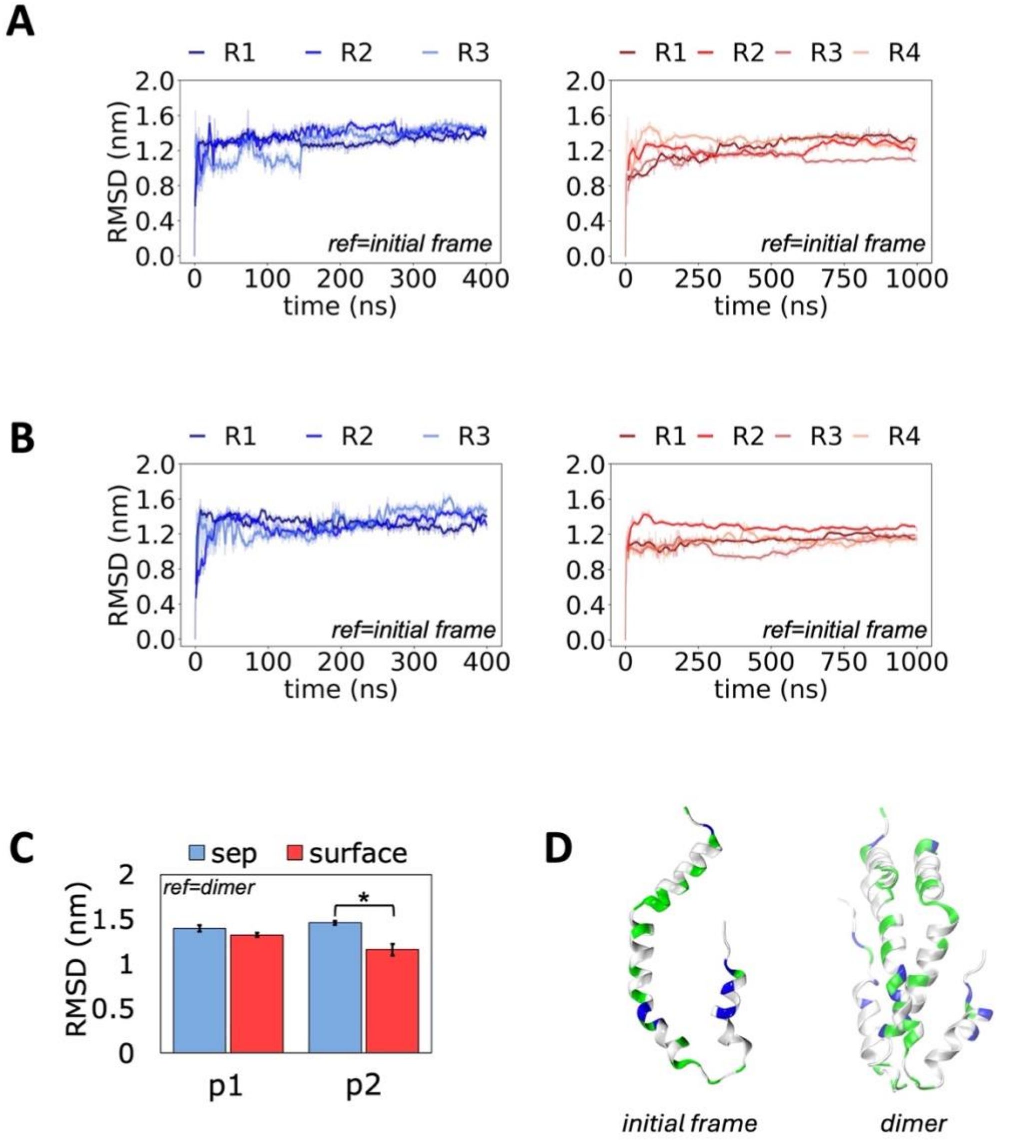


**Figure S2. A)** RMSD of monomer 1 (p1) and **B)** monomer 2 (p2) calculated using the first frame of each protein as the reference in each case. Results for *Sep* model on the left, and *Surface* on the right. **C)** Average RMSD of *Sep* and *Surface* monomers using the *Bound* dimer structure as the reference. **D)** Reference structures used to calculate RMSD. Error bars represent standard error across replicas, and "*" indicates significant difference in means ($p < 0.05$).

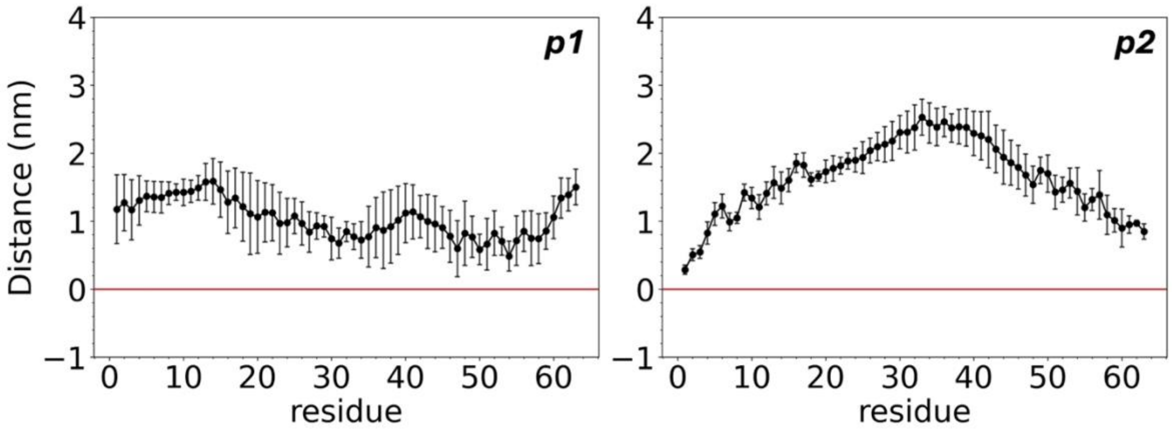


**Figure S3.** Distance of each amino acid from the average position of phosphorus atoms in the contacting membrane leaflet in the *Surface* systems. The average position of phosphorus atoms is indicated with the red line at 0. Error bars represent standard error across replicas.

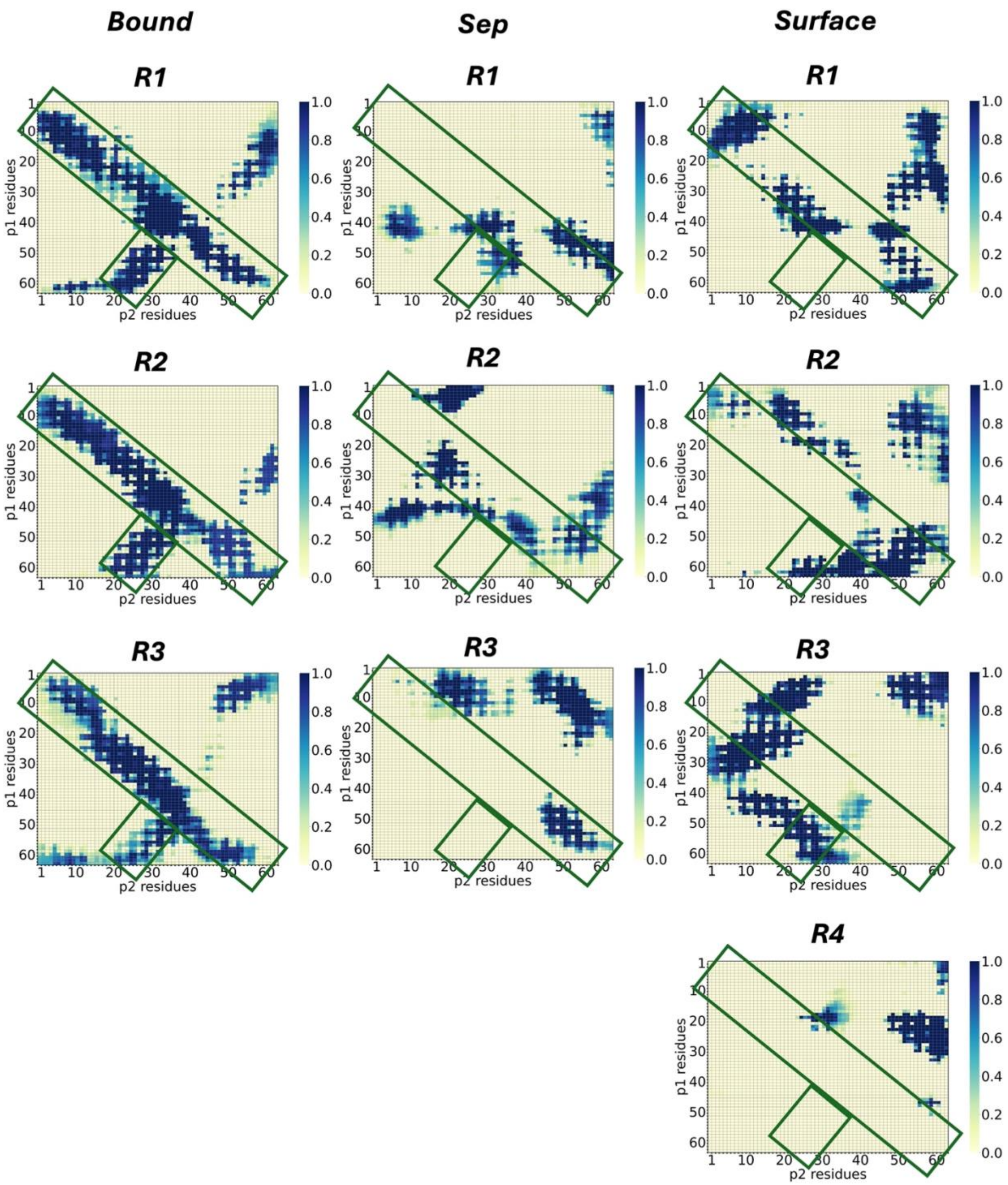


**Figure S4.** Contact maps of residues during the last half of trajectory (200 ns in *Sep* and 500 ns in *Surface* model), based on a cutoff of 14 Å. Green rectangles indicate region populated in all *Bound* model replicas. Top to bottom panels correspond to either replicas 1-3 (*Bound* and *Sep* models) or 1-4 (*Surface* model).

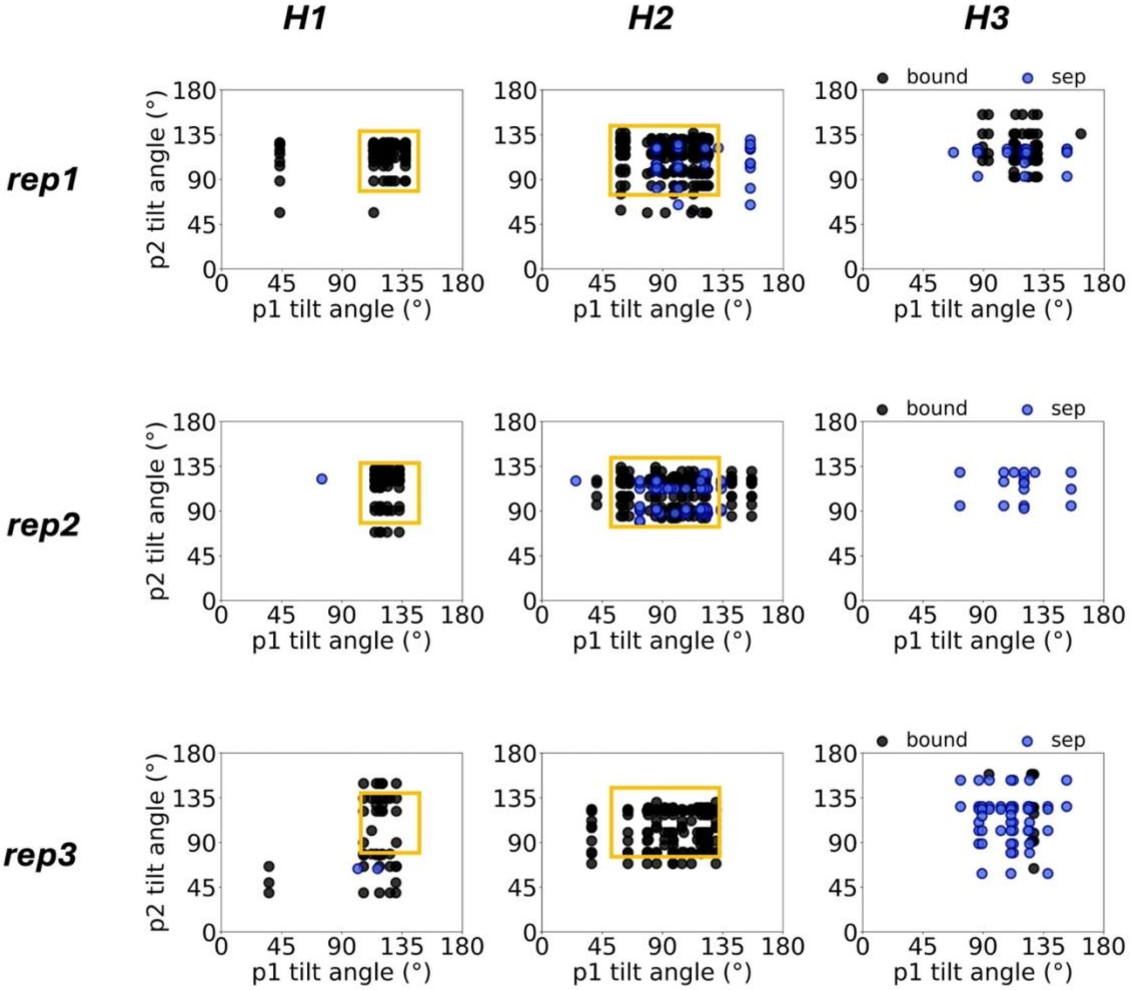


**Figure S5.** Tilt angle conformational landscapes of residues involved in 1-to-1 helix contacts during the last 200 ns of trajectory in the *Sep* model. Results for helices 1, 2 and 3 shown in the left, middle and right columns, and replicas 1, 2 and 3 shown in the top, middle and last rows. Yellow rectangles indicate populated regions conserved across all *Bound* replicas.

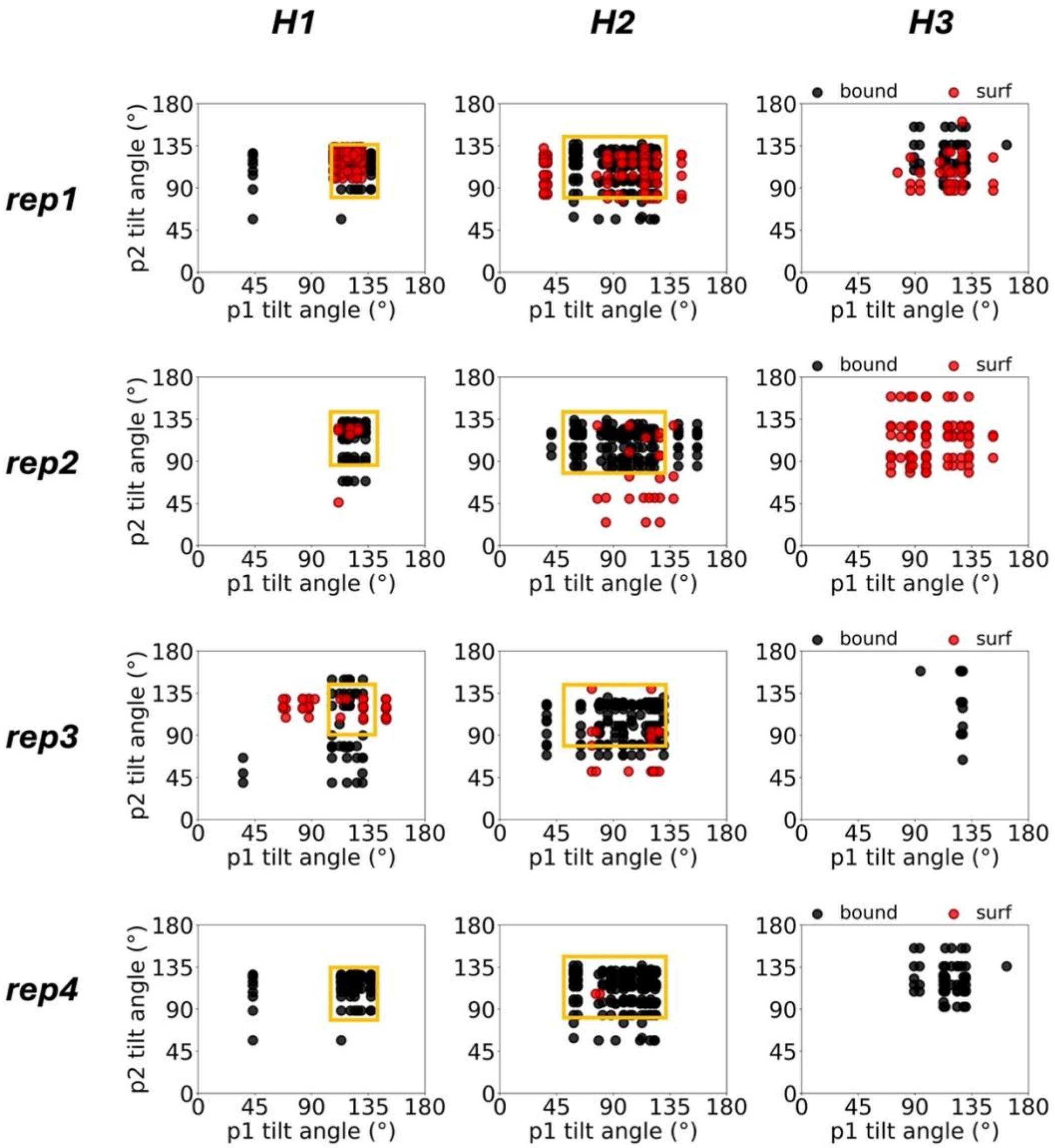


**Figure S6.** Tilt angle conformational landscapes of residues involved in 1-to-1 helix contacts during the last 500 ns of trajectory in the *Surface* model. Results for helices 1, 2 and 3 shown in the left, middle and right columns, and replicas 1, 2, 3 and 4 shown in the corresponding rows. Yellow rectangles indicate populated regions conserved across all *Bound* replicas.

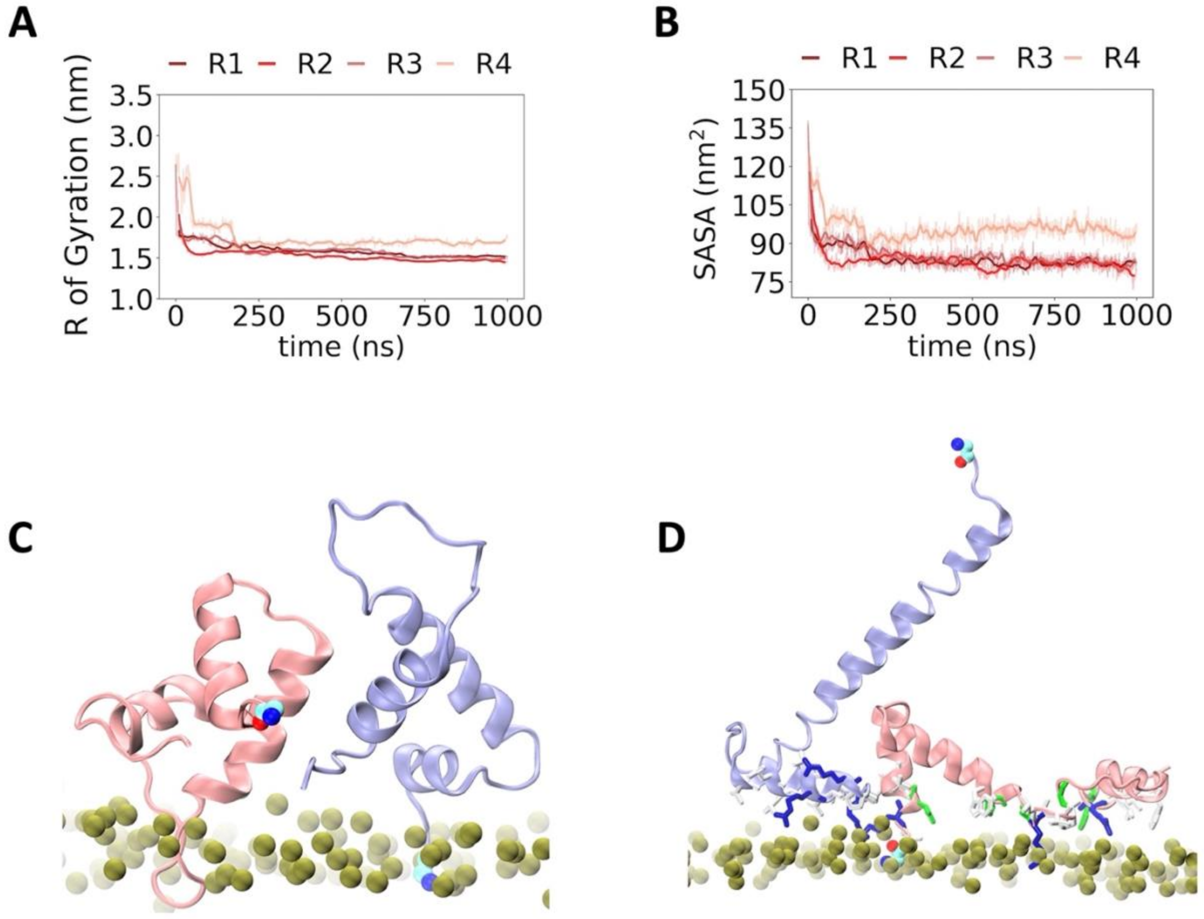


**Figure S7.** Time series of **A)** $R_g$ and **B)** SASA of *Surface* replica 4. **C)** Final conformation, and **D)** 27ns snapshot of the conformation of the dimer on the contacting membrane leaflet in this replica; phosphorus atoms are shown in green for reference. Proteins differentiated with pink (p1) and ice-blue (p2), with the N-terminus end indicated with van der Waal representation. In panel D, the residues within 8 Å of the phosphorus atoms are also shown, with nonpolar residues in white, polar in green, cationic in blue.

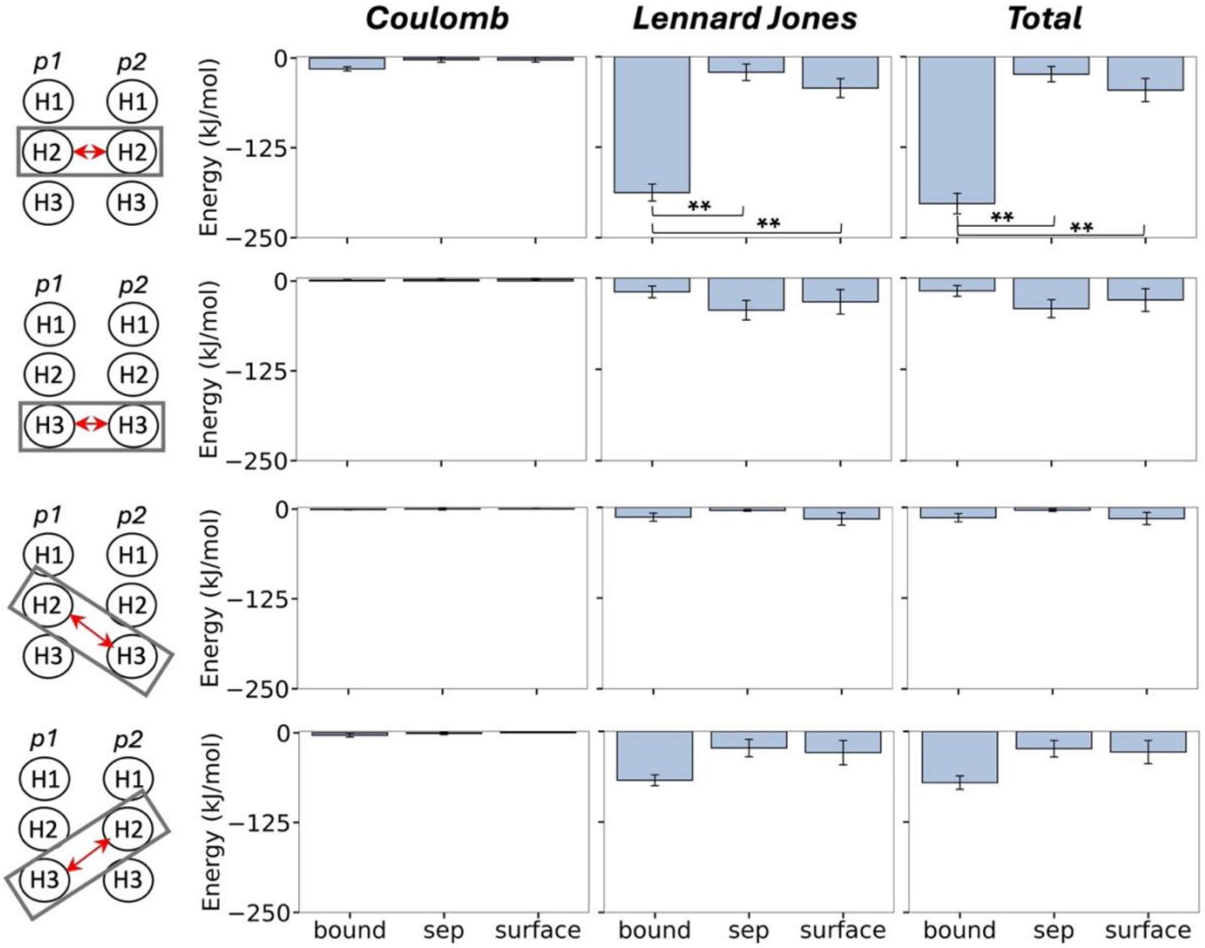


**Figure S8.** Coulomb (electrostatic), Lennard-Jones (hydrophobic), and total interaction energy of interacting monomer helices in each model. Error bars represent standard error across replicas, and "**" indicates significant difference in means ($p < 0.01$). Only non-zero estimates of p1 and p2 helix interaction energies are shown.

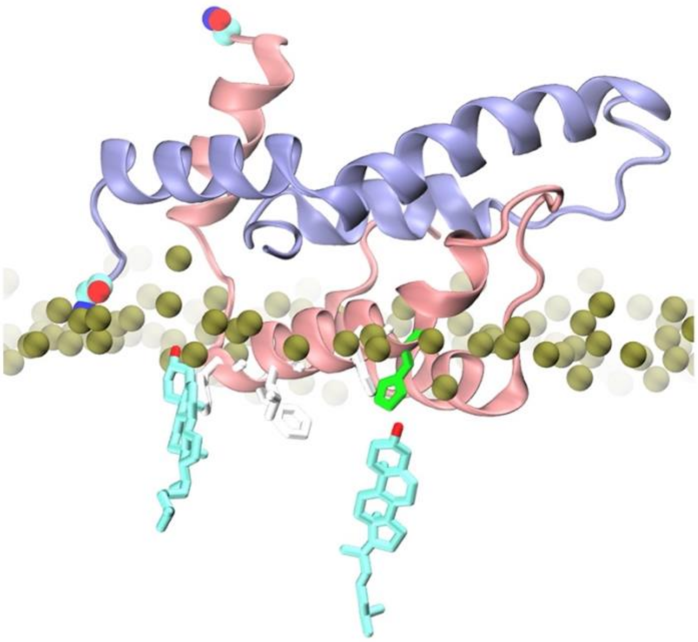

**Figure S9.** Snapshot showing cholesterol contacts with the N-terminus of p2 and H2 and H3 of p1 in the *Surface* model. Lipid phosphorus atoms are shown in green as reference, the rest of the lipid structure, water, and ions are hidden for clarity. Proteins differentiated in pink and ice-blue, with the N-terminus end indicated with van der Waal representation. Non-polar and polar protein residues in white and green respectively.

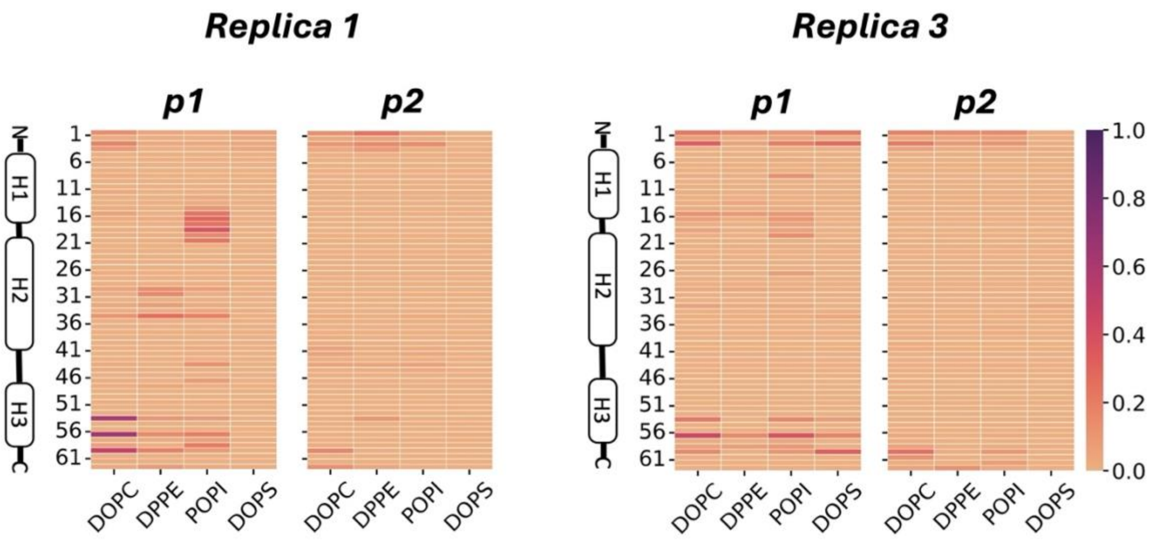


**Figure S10** Frequency of hydrogen bonds between individual residues and lipid species during the last half of trajectory (500 ns) in the *Surface* model. Results shown are for representative *Surface* replicas that form the most accurate dimer contact configuration.

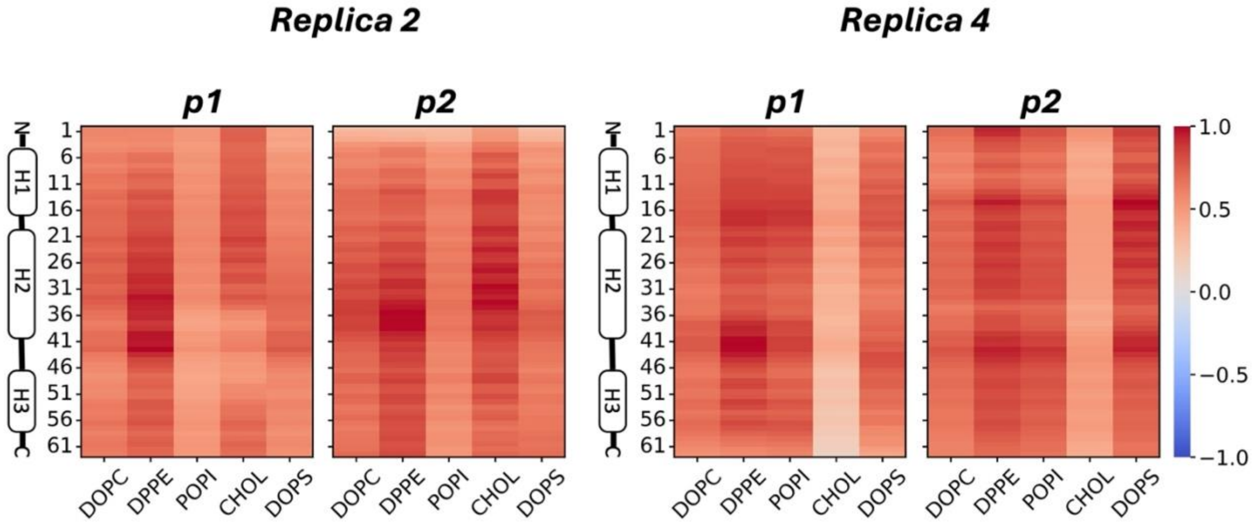


**Figure S11.** Protein-lipid dynamic cross correlation maps for *Surface* replicas 2 and 4 during the entire trajectory. Negative and positive correlations represented in blue and red respectively.
